# Comparative analysis of wild type accessions reveals novel determinants of Arabidopsis seed longevity

**DOI:** 10.1101/2021.09.13.460022

**Authors:** Regina Niñoles, Dolores Planes, Paloma Arjona, Carmen Ruiz-Pastor, Rubén Chazarra, Joan Renard, Eduardo Bueso, Javier Forment, Ramón Serrano, Ilse Kranner, Thomas Roach, José Gadea

**Affiliations:** Instituto de Biología Molecular y Celular de Plantas (IBMCP), Universitat Politècnica de València (UPV)-Consejo Superior de Investigaciones Científicas (CSIC), Ciudad Politécnica de la Innovación (CPI), Ed. 8E, C/Ingeniero Fausto Elio s/n, 46022, Valencia, Spain.; Department of Botany and Center for Molecular Biosciences Innsbruck (CMBI), University of Innsbruck, Innsbruck, Austria.

**Keywords:** Seed, longevity, transcriptomics, natural variation, Arabidopsis, glutathione

## Abstract

Understanding the genetic factors involved in seed longevity is of paramount importance in agricultural and ecological contexts. The polygenic nature of this trait suggests that many of them remain undiscovered. Here, we exploited the contrasting seed longevity found amongst wild type Arabidopsis thaliana accessions to further understand this phenomenon. Concentrations of the antioxidant glutathione were consistently higher in longer-lived than shorter-lived accessions, supporting that redox poise plays a prominent role in seed longevity. However, high seed permeability, normally associated with shorter longevity, is also present in accessions with longer seed longevity. Transcriptome analysis indicated that the detrimental effect on longevity caused by seed coat permeability may be counterbalanced by higher levels of specific mRNAs stored in dry seed, particularly those of heat-shock proteins. Indeed, reverse genetics demonstrated that heat-shock factors HSF1A and 1B contributed to longevity. Furthermore, loss-of-function mutants of RNA-binding proteins, such as the stress-granule zinc-finger protein TZF9, or the spliceosome subunits MOS4 or MAC3A/MAC3B, extended seed longevity, positioning RNA as a novel player in the regulation of seed viability. mRNAs of proteins with putative relevance to longevity were also abundant in shorter-lived accessions, reinforcing the idea that resistance to ageing is determined by multiple factors.

## INTRODUCTION

Understanding why some seeds survive for years or centuries while others die within months is a paramount question in seed science. Further to genetics, environmental cues perceived by the mother plant during seed maturation (He et al, 2014; Nagel et al, 2015), harvesting practices (Probert et al, 2007), postharvest treatments, such as seed priming (Wang et al, 2018), and storage conditions (Walters et al, 2004), all influence seed longevity. Regarding the genetic basis, QTL and genome-wide association studies suggest involvement of many loci, underlining the highly polygenic nature of seed longevity (Agacka-Modoch et al, 2015, Nagel et al, 2015, Renard et al, 2020b).

The desiccated state of orthodox seeds restricts metabolism (Buitink and Leprince 2008), which extends their life-span, or ‘longevity’, well beyond what is capable for desiccation intolerant ‘recalcitrant’ seeds (Angelovici et al. 2010). Nonetheless, a loss of redox homeostasis, presumably due to excess ROS production, coincides with viability loss in both recalcitrant and orthodox seeds (Kranner et al., 2006; Roach et al., 2010). As for all desiccation tolerant organisms, life in a metabolically-inert state entails accumulation of oxidative damage, eventually resulting in cell death. A plethora of metabolites, including antioxidants like glutathione and tocochromanols (Kranner et al, 2006; Nagel et al, 2015) (Sattler et al, 2004), and various seed storage (Nguyen et al, 2015) or heat-shock proteins (Tejedor-Cano et al, 2010; Bissoli et al, 2012), help reduce damage while in the desiccated state (Kranner and Birtic, 2005)(Bailly 2004)(Sano et al., 2016). Glutathione has a dominant role in maintaining the cellular environment in a reduced redox state, necessary for efficient metabolism, redox signalling and protection from oxidative damage. Seed ageing is associated with a decrease in GSH and increase in GSSG concentration, in other words, seeds oxidise during ageing as they lose vigour and viability. Based on molar concentrations of GSH and GSSG, seed cellular redox state can be quantified via the Nernst equation (E_GSSG_/_2GSH_), which has a sigmoidal relationship with seed lot viability (Kranner et al, 2006). The maternal-derived seed coat is a partial barrier to oxygen, minimizing oxidation of cellular components (Renard et al., 2021). Mutants affected in seed coat development (Leon-Kloosterziel et al, 1994) or in the composition of the integument lipidic layers, have high seed coat permeability and low viability, suggesting that both factors are inversely correlated (Renard et al, 2020a; Demonsais et al, 2020).

During germination, pre-existing mRNAs, stored in mature dry seeds, are used for protein synthesis. More than 12,000 different types of mRNAs are stored in the dry seed of Arabidopsis thaliana. They include mRNAs needed for the repair of cellular damage and for initiation of the germination process, together with others that persist from seed development (Nakabayashi et al, 2005). Dry seed transcriptomes of the Col-0 and Cvi accessions of Arabidopsis resemble one another, but a fraction of stored mRNAs is accession-specific (Kimura and Nambara, 2010). It is plausible that the nature and abundance of this unique pattern of stored mRNAs between accessions could explain the diversity observed in seed properties. In this regard, Col-0 and Est-1 accessions, which greatly vary in longevity, also differ in a substantial fraction of their dry seed transcriptomes (Sano et al, 2017). Furthermore, proteomes of wheat genotypes with contrasting seed longevities revealed differences in abundance of one-sixth of all detected proteins (Chen et al, 2018). If there are specific proteins within this list conferring or contributing to longevity is unknown.

Reverse genetics studies using systematic or candidate genes approaches have revealed many cellular components involved in seed longevity in Arabidopsis (Bueso et al, 2014, 2016). Co-expression network analysis identified a list of novel genes and processes likely affecting viability (Righetti et al, 2015); recently, the success of this approach was demonstrated, revealing a novel role for auxin signalling in seed longevity (Pellizzaro et al, 2020). We recently used large-scale phenotyping, coupled with genome-wide association studies and reverse genetics, to identify novel genes involved in seed deterioration (Renard et al, 2020b). Comparative analyses of natural accessions with contrasting tolerance is an alternative strategy to elucidate molecular factors and mechanisms involved in stress tolerance (Veen et al, 2016; Kusunoki et al, 2017). Moreover, natural accessions are an adequate material to investigate in which ways adaptations to different traits are tuned to provide an optimum response for the plant. However, this approach has yet to be applied to seed longevity in a systematic way.

In this study, to investigate attributes of seed longevity from multiple angles, we compared seed coat permeability, dry seed mRNA composition and glutathione contents, in wild type accessions of Arabidopsis with different relative longevity, referred to as “long-lived” and “short-lived”. A positive correlation of glutathione contents at seed maturity with seed life span supports that seed oxidation accompanies viability loss, regardless of ageing rate. The importance of heat-shock proteins in resisting ageing was supported in this study, and the transcription factors HSF1A and HSF1B appear to be important regulators in seeds. Moreover, by assaying longevity in T-DNA insertional mutants of genes of the splicing machinery or mRNA packing regulation, we revealed an unexplored role of RNA processing in seed longevity, positioning RNA as a key player in the regulation of seed viability. However, we also highlight the multifactorial character of seed longevity, and reveal potential redundancy in some putative processes employed by seeds to resist ageing when plants are grown under controlled conditions.

## MATERIALS AND METHODS

### Plant Material and growth conditions

Arabidopsis thaliana ecotypes were obtained from the Nottingham Arabidopsis Stock Centre (NASC) (reference N76309). All Ecotypes were grown simultaneously to obtain fresh seeds in homogeneous conditions (16 hr light/8 hr dark, at 22°C and 70–75% RH). For T-DNA insertion lines, the tzf9 line was kindly donated by Dr. Justin Lee (Maldonado-Bonilla et al, 2014); the mac3a/mac3b and mos4-1 lines were obtained from NASC, and hsf1/3 was obtained from Dr. F. Schöffl’s laboratory (Lohmann et al, 2004). Seeds were harvested from Arabidopsis plants grown under greenhouse conditions (16 hr light/8 hr dark, at 23±2 °C and 70 ± 5% relative humidity) in pots containing a 1:2 vermiculite:soil mixture.

Seeds were surface sterilized with 70% ethanol containing 0.1% Triton X-100 for 15 min followed by rinsing three times with sterile water. Seeds were stratified for 3 days at 4 °C. Germination was carried out on plates containing Murashige and Skoog (MS) salts with 1% (w/v) sucrose, 10 mM 2-(N-morpholino) ethanesulfonic acid and 0,9% (w/v) agar, pH was adjusted to 5.7.

### Seed ageing assays

Ambient ageing was performed by storing dry seeds at ambient temperature and humidity (20-25°C, 40-60% RH) for 18 or 30 months, as indicated in each case. The Controlled-Deterioration Treatment (CDT) consisted of maintaining seeds during 14 or 21 days at 37°C or 40°C, as indicated in each case, in an atmosphere of 75% HR above a saturated NaCl solution.

### Seed coat permeability test

Triphenyltetrazolium reduction assay was used as described in Molina et al, 2008. Briefly, seeds were incubated in the dark in 1% (w/v) tetrazolium red at 30 °C for 36 hr. Then, formazan was extracted and quantified as the absorbance at 485 nm of 1ml sample. Values shown in figures are means of three biological replicates. Each replicate contains 20 mg of seeds.

### Glutathione thiol and disulphide measurements

Seeds (15-20 mg / replicate) were freeze-dried for 5 d before grinding with two 3 mm quartz beads at 30 Hz for 2 min at -80 °C. 1 mL of ice-cold 0.1 M HCl and polyvinylpolypyrrolidone equal to seed weight was added before extracting at 30 Hz for 2 min. The extract was centrifuged at 20,000 × g for 10 min at 4 °C and 700 µL of the supernatant was transferred, avoiding the lipid phase, for re-centrifugation as before. LMW thiols and disulphides were analysed by HPLC modified from (Kranner, 1998). Briefly, for the determination of total LMW thiols and disulphides (e.g. GSH+GSSG), 120 µL of the supernatant was pH-adjusted to 8.0 with 180µL of 200 mM bicine buffer, reduced with dithiothreitol (DTT) for 1 h, before labelling thiol groups with monobromobimane (mBBr) for 15 min. To quantify disulphides only (e.g. GSSG), thiols (e.g. GSH) in 400 µL of supernatant were blocked for 15 min with N-ethylmaleimide (NEM), which was added just before pH-adjustment with 600 µL bicine buffer. Excess NEM was removed with an equal volume of toluene four times. Disulphides in 300 µL of NEM-treated extract were reduced with DTT and labelled with mBBr, as before. mBBR-labelled compounds were separated by reversed-phase HPLC, using an Agilent 1100 HPLC system (Agilent Technologies, Santa Clara, CA, USA), on a ChromBudget 120-5-C18 column (5.0 µm, BISCHOFF GmbH, Leonberg, Germany), and detected by a fluorescence detector (ex: 380 nm, em: 480 nm). Amount of each disulphide was calculated by halving the number of thiol equivalents measured (e.g. 1 mol GSSG = 2 mol GSH), and amount of each LMW thiol was calculated by subtracting the amount of LMW thiol equivalents of each disulphide from total amounts of LMW thiol equivalents of both disulphides and thiols, e.g. GSH = (mol GSH+ (mol GSSG x 2)) - (mol GSSG x 2).

The half-cell reduction potential (E_hc_) for each thiol-disulphide couple was calculated according to the Nernst equation (Kranner et al., 2006; Schafer and Buettner, 2001):

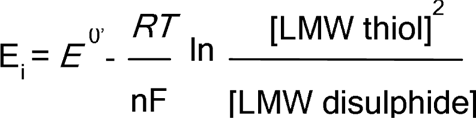

where R is the gas constant (8.314 J K^-1^ mol^-1^ ); T, temperature in K^-1^; n, number of transferred electrons (=2); F, Faraday constant (9.6485 x 10 C mol ); E , standard half-cell reduction potential of a thiol-disulphide redox couple at an assumed cellular pH of 7.3 (E^0’^ _GSSG/2GSH_ = -258 mV). E_i_ is the half-cell reduction potential of an individual redox couple i, and [reduced species]_i_ is the concentration of the reduced species in that redox pair. The molar concentrations of LMW thiols and disulphides were calculated based on seed water content.

### RNA extraction and qRT-PCRs

Extraction of total RNA from dry seeds was performed according to Oñate-Sánchez and Vicente-Carbajosa (2008). DNA removal was performed with DNAse kit (E.Z.N.A). About 1 μg RNA was reverse-transcribed using the Maxima first-strand cDNA synthesis kit (Thermo Fisher Scientific) according to the manufacturer’s instructions. qRT-PCR was performed with the PyroTaq EvaGreen qPCR Mix Plus (ROX; Cultek S.L.U., Madrid, Spain) in a total volume of 20 μl using an Applied Biosystems 7500 Real-Time PCR System (Thermo Fisher Scientific). Data are the mean of three replicates. PCR amplification specificity was confirmed with a heat-dissociation curve (from 60 to 95 °C). The internal standard used was ASAR1 (AT4G02080), specifically selected for its stable expression in seeds (Bas et al, 2012). Relative mRNA abundance was calculated using the comparative ΔCt method (Pfaffl, 2001). Primers for qRT-PCR are listed in Supplemental table 1.

### Transcriptomic assays and RNA data analysis

RNA was extracted from dry seeds as mentioned before. Twenty-million paired-ends 50pb reads per library (40 million in total) were sequenced. Two or three replicates were used per accession. After adaptor removal and low-quality trimming of raw reads with cutadapt (Martin, 2011), clean reads were quality assessed with FastQC (Andrews, 2010) and mapped to the TAIR10 Arabidopsis thaliana genome using HISAT2 (Kim et al, 2015). Gene counts were then obtained with htseq-count (Anders et al, 2015) and used for differential expression analysis with DESeq2 (Love et al, 2014). For PCA and clustering analysis, average expression (read-per-kilobase-million reads, RPKM) was used per accession, displaying only genes with standard deviation higher than 500 between accessions. UPGMA was used with average linkage and Pearson correlation. Functional analysis was performed using AgriGOv2 (Tian et al, 2017). Redundant GO terms were removed and remaining GO terms were visualised using ReviGO (Supek et al, 2011). For differential splicing analysis, quantification of AtRTD2 transcript isoforms (Zhang et al, 2016) was done with salmon in mapping-based mode (Patro et al, 2017) and used for alternative splicing analysis with IsoformSwitchAnalyzeR (Vitting-Seerup ert al, 2019). Data have been submitted to GEO repository under the accession number GSE179008.

## RESULTS

### Arabidopsis accessions differ in seed longevity

For the present study 30 accessions were selected for further analysis, 14 “long-lived” accessions with higher total germination (TG) and 16 “short-lived” ones with lower TG than Col-0 after 18 months of ambient ageing (Figure 1a). After 30 months, seed longevity was re-assayed in triplicate for 14 of these accessions (8 with TG> and 5 with TG< than Col-0) (Figure 1b). The same 14 genotypes were also aged much more rapidly using CDT at 37°C and 75% RH. Total germination of these 14 genotypes after these contrasting ageing regimes significantly correlated (R^2^ =0.911; Pearson R = 0.955, Figure 1c). As all plants were grown under the same controlled environmental conditions during seed development, these results suggest that genetic differences affecting seed traits were responsible for the distinct longevity responses.

**Figure 1.**
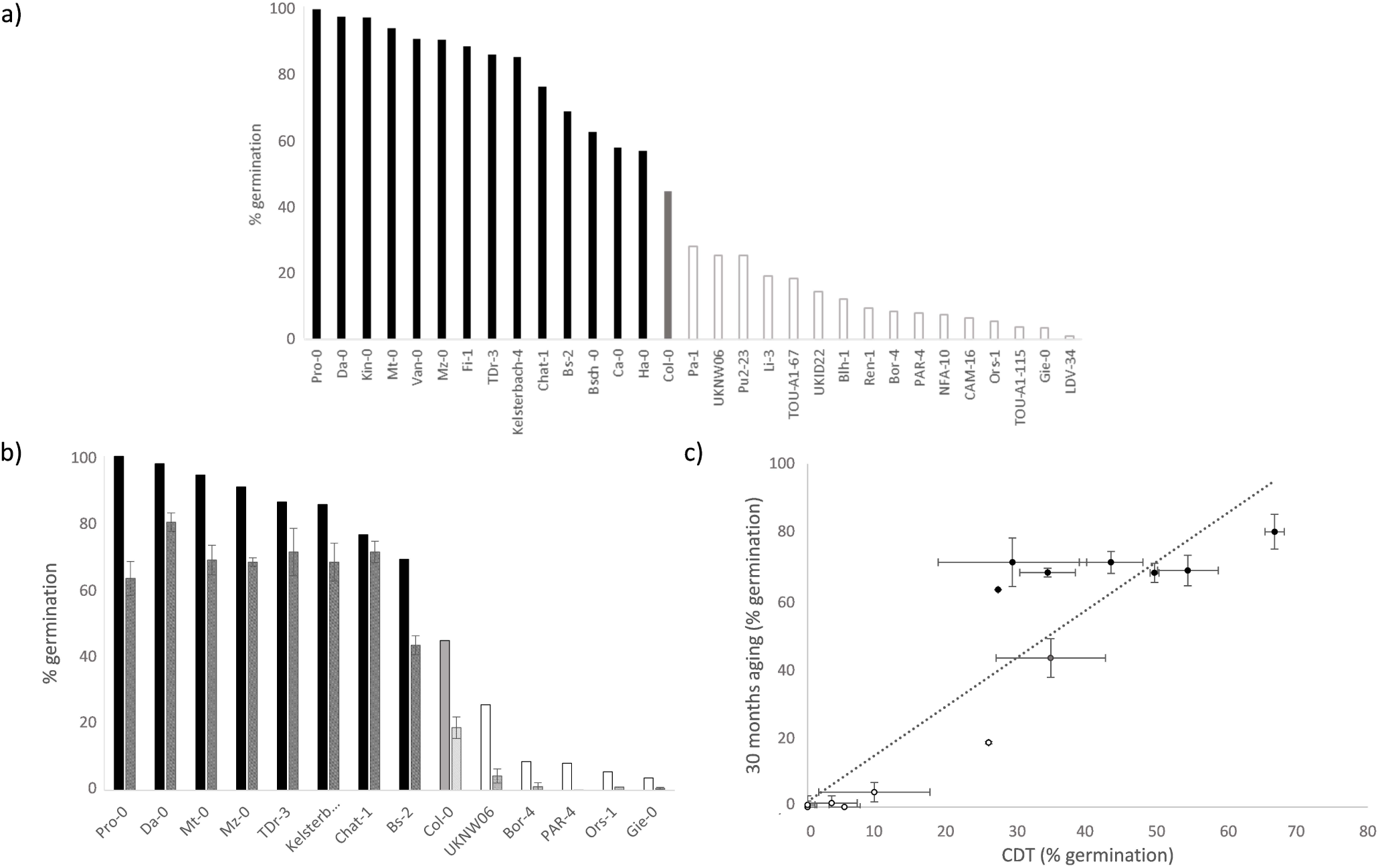
Phenotypic response to seed deterioration in Arabidopsis thaliana accessions. a) Percentage of total germination of long-lived (black bars) and short-lived (white bars) accessions after 18 months of ambient ageing according to Renard et al, 2020. Col-0 has intermediate seed longevity and is included for reference (grey bar) b) Percentage of germination of 8 long-lived and 5 short-lived accessions after 30 months of ambient ageing (grey bars). Results of triplicate assays are compared to data from Figure 1a. c) Correlation of data obtained by 30-months of ambient ageing and controlled-deterioration for extreme accessions. Black solid circles: long-lived accessions; open circles: short-lived accessions; grey circle: Col-0.

### High seed coat permeability and low longevity does not always correlate

The tetrazolium penetration assay conducted on seed batches showing 100% total germination is a standard method to quantify seed coat permeability (Molina et al, 2008). High seed coat permeability is a feature of flavonoid-deficient transparent-testa (tt) mutants, which tend to have shortened seed longevity (Debeaujon et al., 2000). To test if the inverse association of seed coat permeability with seed longevity applies to wild type accessions, we assayed tetrazolium permeability in 30 accessions with contrasting seed longevity. No correlation with seed longevity was observed (Figure 2b, r= -0,28). In agreement with flavonoid-deficient tt mutants, very high permeability (A_485_>0.6) was associated with shortened longevity, although accessions Bs-2, Da-0, Bsch-0 and TDr-3 are long-lived despite being quite permeable (A_485_>0.5). On the other hand, although very low permeability (A_485_<0.2) was generally found in long lived accessions, there were exceptions to the rule, e.g. the short-lived PAR-4 had very low permeability. Moreover, no clear trend was observed in the intermediate range (A_485_=0.2 to 0.5), with accessions of contrasting longevity (Figure 2a). In summary, these results indicate that other mechanisms likely exist that compensate highly permeable seed coats in accessions that are also relatively long lived. Similarly, other seed properties may be responsible for the short lived phenotype of accessions with less permeable seed coats.

**Figure 2.**
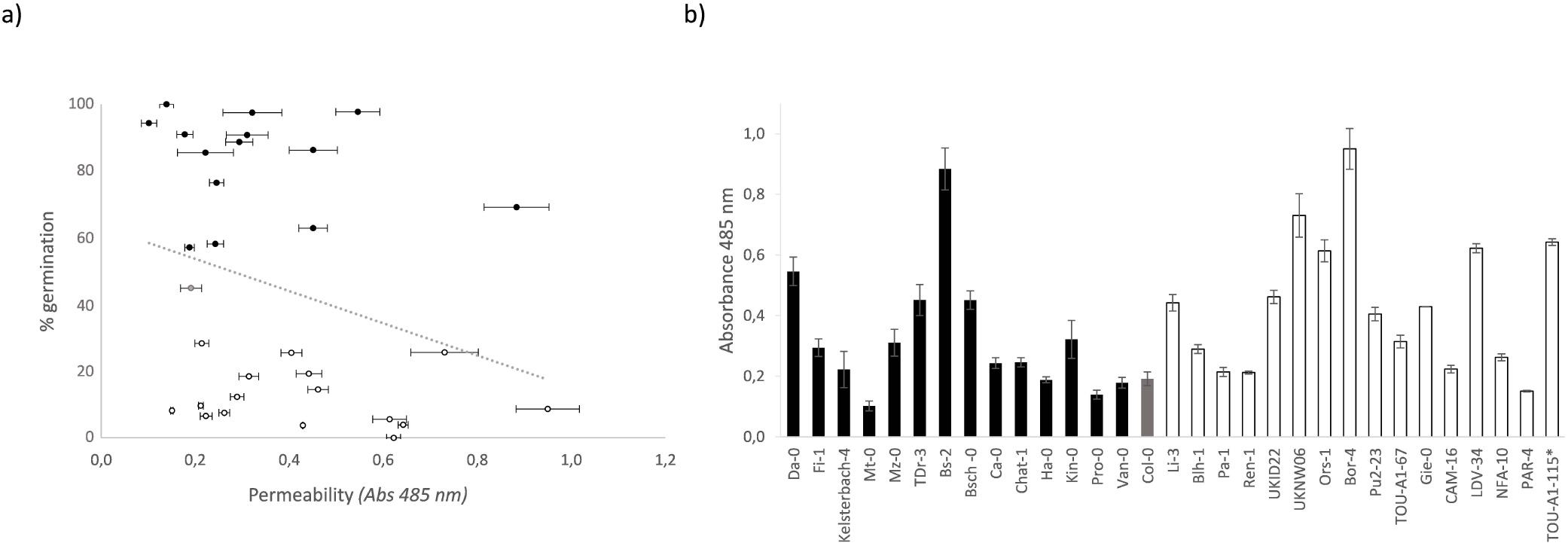
a) Tetrazolium assay in long-lived (black bars) and short-lived (white bars) accessions. Col-0 has an intermediate response for seed longevity (grey bar). b) Correlation between total germination after 18-months of ambient ageing and permeability data according to the tetrazolium assay in Figure 2a.

### Stored mRNAs composition differs between high and shorter-lived accessions

Next, we conducted a transcriptome analysis to compare stored mRNAs composition in long- and short-lived accessions. In total, six accessions with long-lived seeds (Da-0, Pro-0, TDr-3, Chat-1, Kelsterbach-4 and Fi-1) and six with short -lived seeds (Ors-1, Bor-0, PAR-4, UKNW06-386, CAM-16 and Ren-1), relative to Col-0, were analysed. Thirty libraries were prepared to generate around 20,000 high-quality 50 bp single-end reads per library (Supplemental Table 2). Around 12,000 genes were expressed in dry seeds and transcriptomes of different accessions resembled one another, as previously reported (Nakabayashi et al, 2005): around 80% of the genes identified in the seed were expressed in all accessions (see Table 1), and only 737 genes were accession-specific. However, for all accessions, a non-negligible fraction of the dry seed transcriptome (10-28%, depending on the accession, see Table 1, was differentially expressed as compared to Col-0, indicating differences in transcript abundance between them.

**Table 1.**
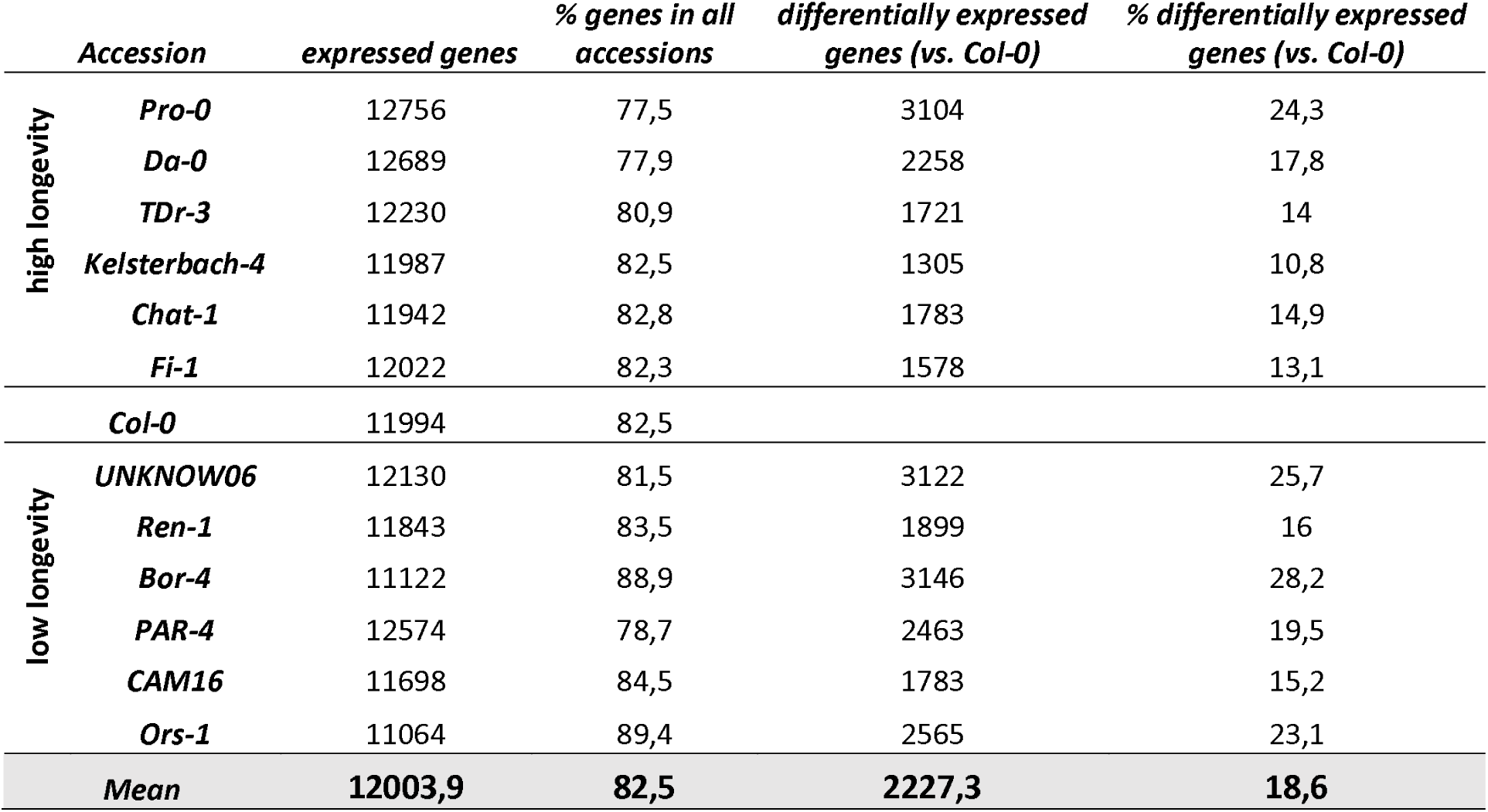
mRNAs present in dry seeds of wild type accessions of Arabidopsis, and differentially expressed genes compared to Col-0. Expressed genes: Number of genes representing at least two replicates with RPKM>1 (reads per kilobase per million reads); % of genes in all accessions: percentage of expressed genes with transcripts in all accessions (including Col-0); differentially-expressed genes (vs. Col-0): differentially expressed genes with FDR adjusted p-value <0.05 and log_2_FoldChange < -1 or > 1; % differentially expressed genes (vs. Col-0): percentage of differentially expressed genes vs. Col-0 from the total number of genes expressed in each accession.

Principal component analysis (PCA) was performed to estimate the relationship between dry seed transcriptomes and longevity. As shown in Figure 3, the first PC (PC1) explained most of the total variation observed. A trend was observed with PC1 separating samples according to longevity, the long-lived accessions mapping to the top-left quadrant of the plot, and the short-lived accessions distributed along the middle- and bottom-right quadrants. This indicates that differences in abundance of stored mRNAs between accessions could help explain the observed differences in seed longevity. However, separation by PCA was not perfect, and a cluster of long- and short-lived accessions grouped quite close to the intermediate Col-0, indicating that, at least for those ones, transcript provisions are similar, and suggesting that there is not a common characteristic in the transcriptome that explains the longevity phenotypes for all accessions. Only two genes were found of which the expression levels correlated with longevity (see Supplemental Table 3). The glycosyltransferase IAGLU (AT4G15550), which glycosylates kaempferol (Meßner et al, 2003), was more abundant in the six long-lived accessions and less abundant in five of the short-lived accessions. On the contrary, the phytochrome interacting factor 3 (PIF3, AT1G09530), a transcription factor transducing phytochrome signals (Monte et al, 2004), was less abundant in the six short-lived accessions analysed. Although a contribution of any of these two genes to longevity cannot be discarded, the fact that a positive trend between expression and longevity was not observed for any other gene when considering all the accessions together (Supplemental Table 3) reduces the possibility to find a common feature in the transcriptome that explains the longevity phenotypes.

**Figure 3.**
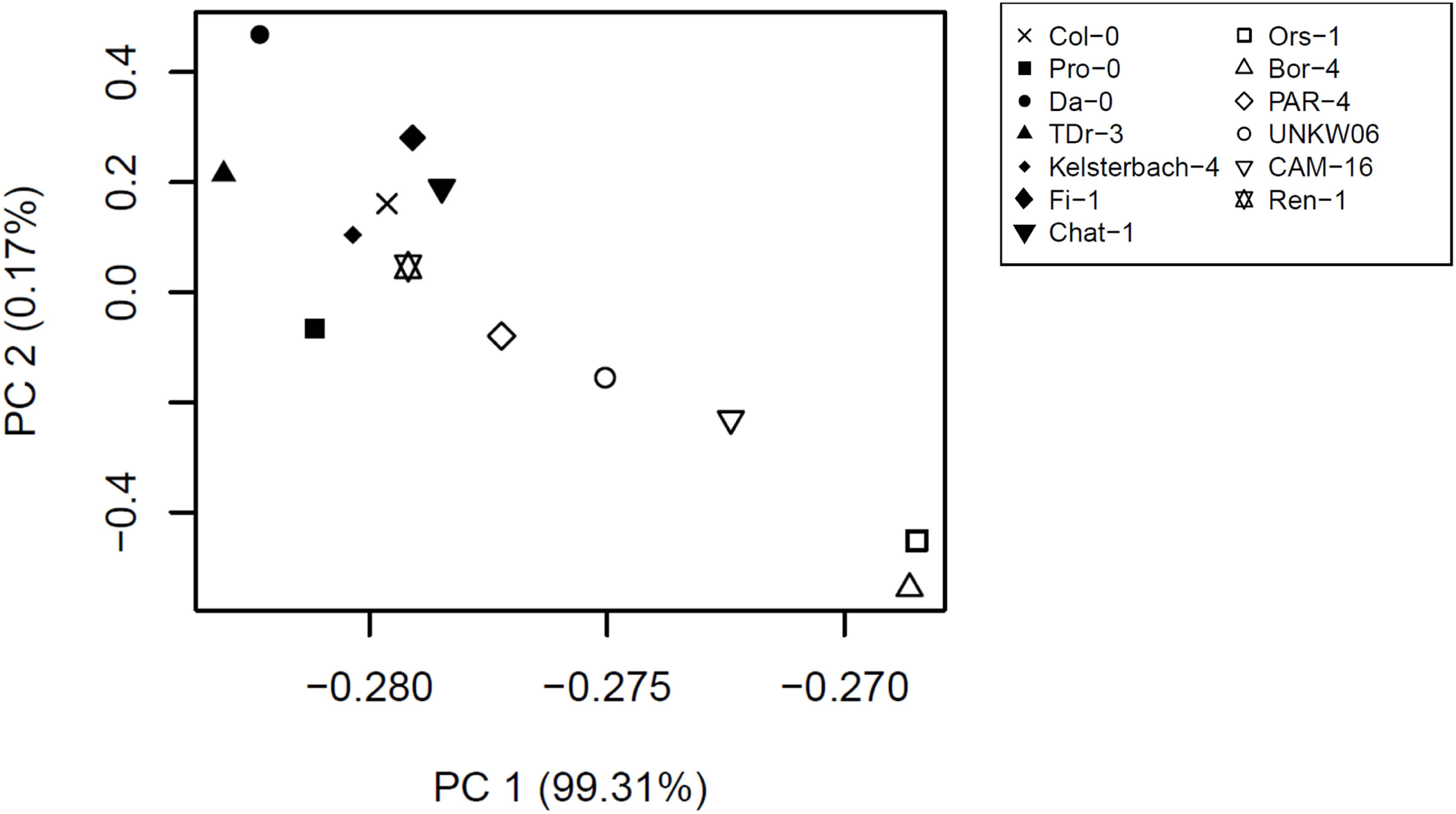
Principal component analysis of transcriptome data of long-lived (black symbols) and short-lived (white-symbols) accessions. Col-0, with intermediate seed longevity, is included for reference (grey). Average of read counts was used for every accession. Most variable genes (those showing a standard deviation of count data in all accessions higher than 500) were included in the calculation.

For some of the accessions, however, the separation along PC1 was pronounced, and as indicated, this separation was correlated with longevity, suggesting that comparison of stored mRNAs could provide insights into new components involved in seed longevity.

### Both short- and long-lived seeds have stored mRNAs important for seed longevity

To highlight genes that could explain differences in seed longevity, the long-lived Da-0 accession was compared to the short-lived Ors-1 and Bor-4 (Supplemental Table 4). They mapped apart in the PCA plot, with Col-0 in an intermediate position, indicating that they differ considerably in their transcriptome profiles. Cluster analysis highlighted groups of genes with different expression profiles among those accessions. As shown in Figure 4a, clusters 1 and 2 include transcripts more abundant in the long-lived Da-0 than in the short-lived Ors-1 and Bor-4 accessions. Functional analysis using gene ontologies (GO) indicated that these clusters are enriched in genes belonging to five groups of categories (see Figure 4b and Supplemental Table 5). Group 1 categories are related to RNA processing and translational initiation. Genes presenting this pattern of expression include a substantial number of RNA-binding proteins, including the polyA-binding protein PAB2, helicases and splicing factors such as PRP8, SMP2, RBM25, or the components of the MOS4-associated complex MOS4 or MAC3B, as well as translational initiation factors, such as SUI1, IF2/IF5 and several subunits of eIF4 and eIF3. This group also includes the category “glutathione metabolic process”. Group 2 includes categories involved in gene regulation, including “regulation of seed germination”, or “regulation of circadian rhythm”, and includes ERF12, a negative regulator of seed dormancy, repressing the DELAY OF GERMINATION 1 gene, a dormancy master regulator (Li et al, 2019). It also includes the DREB2A-interactor RCD1, involved in oxidative stress responses, the ethylene response factor ERF72, the tocopherol biosynthesis gene VTE1, whose mutation decreases seed longevity (Sattler et al, 2004), the SNRK2.3 kinase, involved in seed development (Nakashima et al, 2009), and some circadian rhythm proteins such as TIME FOR COFFEE or REVEILLE 4. Group 3 includes the category “protein targeting to chloroplasts”, including the import proteins TIC40, TOC159, TOC132 or ALBINO3, important for chloroplast biogenesis. Group 4 includes categories related to seed development, including proteins involved in meristem initiation such as OBE1, OBE2 and TPR1. A higher amount of these transcripts could guarantee a better performance during germination of aged seeds, resulting in enhanced seed longevity. Finally, group 5 includes categories such as “response to oxidative stress” or “response to heat” as clearly enriched. Proteins in those categories include 24 heat-shock proteins or chaperone-like proteins, belonging mainly to the HSP20 subfamily, but also containing some HSP70, HSP90 and HSP100 members, and the heat-shock factor HSFA1E. In the case of the dry seed-specific AT4G27670 (encoding a small chloroplast HSP) and AT4G10250 (encoding an endomembrane-localized HSP20), the abundance in the long-lived accession Da-0 as compared to the short-lived Ors-1 or Bor-4 is 72-fold and 56-fold, respectively. This category also includes the transcription factor DREB2A, which regulates the expression of HSPs via the heat-shock factor 3 (Schramm et al, 2008), as well as the catalase CAT2.

**Figure 4.**
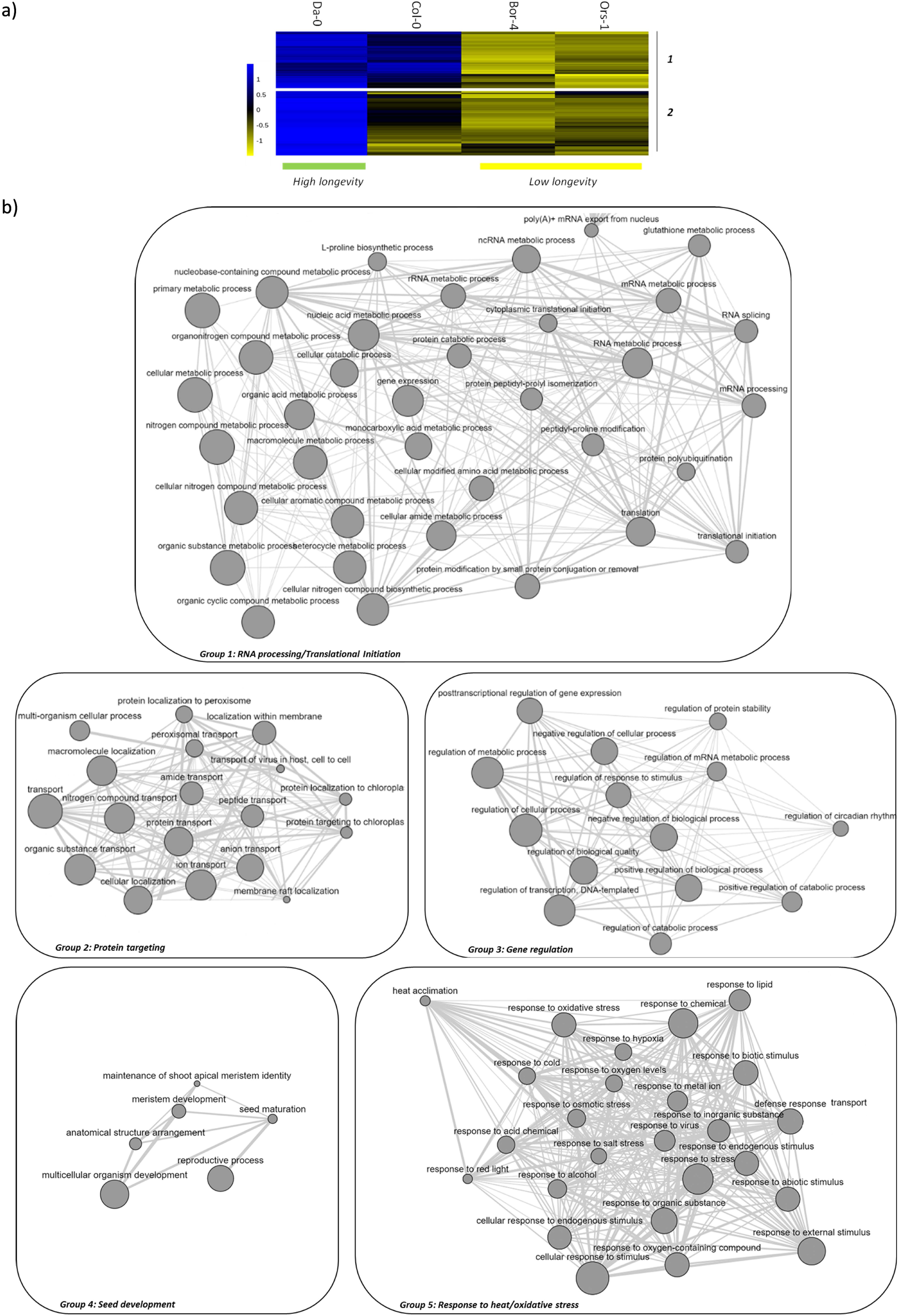
Unsupervised UPGMA clustering analysis (average linkage) over expression data of the long-lived accession Da-0 versus the short-lived accessions Ors-1 and Bor-4. Col-0, with intermediate seed longevity, is included for reference. a) Heatmap showing clusters (1 and 2) containing genes more expressed in Da-0, or in Da-0 and Col-0. b) Gene ontology (GO) analysis (biological process) of the genes belonging to clusters 1 and 2 in Figure 4a. GO analysis was performed using AgriGOv2 (Tian et al, 2017). Redundant GO terms were removed and remaining terms were visualised in the network graph according to ReviGO (Supek et al, 2011).

**Figure 4.**
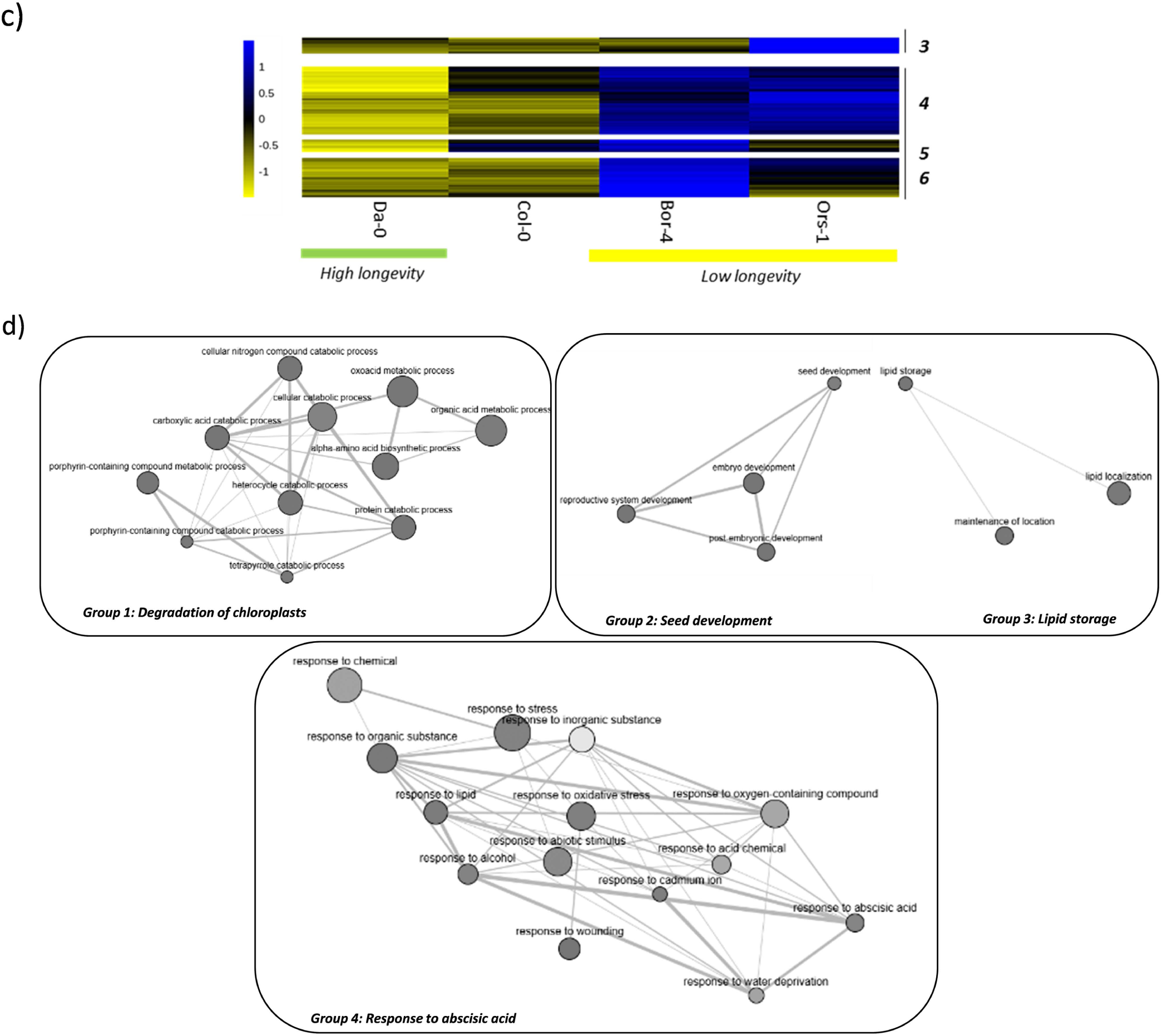
(continued). c) Heatmap showing clusters (3 to 6) containing genes more expressed in Ors-1, Bor-4 or both, than in Da-0 and Col-0. d) Gene ontology (GO) analysis (biological process) of the genes belonging to clusters 3 to 6 in Figure 4c. GO analysis was performed using AgriGOv2 (Tian et al, 2017). Redundant GO terms were removed and remaining terms were visualised in the network graph according to ReviGO (Supek et al, 2011).

Conversely, clusters 3 to 6 (Figure 4c) included transcripts that were more abundant in the short-lived accessions Ors-1, Bor-4 or both. Among the categories enriched in these clusters of genes, four groups could be distinguished (Figure 4d and Supplemental Table 5). Unexpectedly, those categories include genes whose expression has also been associated with increased seed longevity. Group1 includes categories related to the degradation of chloroplasts, with genes such as NYC1 or NYE1, both important for seed maturation and storability (Nakajima et al, 2012; Li et al, 2017). Groups 2 and 3 refer to categories related to “seed development” and “lipid storage”, including genes coding for oleosins, which have been also positively linked with seed longevity (Shao et al, 2019). Finally, in group 4, the category “response to abscisic acid” marked higher abundance of genes induced by this hormone, including the transcription factor ABI5, a key regulator of the transcriptional programme leading to seed desiccation and longevity (Zinsmeister et al, 2016), and 11 late-embryogenesis-abundant (LEA) proteins, induced upon seed maturation drying. Members of the LEA family have also been reported to be beneficial for seed longevity (Hundertmark et al, 2011). Overall, these results indicate that both long-lived and short-lived seeds accumulate different transcripts of proteins that have previously been associated with increased longevity, although this accumulation does not prevent rapid deterioration in short-lived seeds.

Clustering analysis including additional accessions (the long-lived accessions Pro-0 and TDr-3, and the short-lived accessions UKNW06 and CAM-16) (see Supplemental Figure 1) indicates that the genes more expressed in the long-lived ones (cluster 1ext) belong to most of the categories already highlighted by Da-0, (Supplemental Table 5). Remarkably, despite being short-lived, UKNW06 accumulates transcripts more abundant in the long-lived accessions. However, the transcripts accumulated in Bor-4 and Ors-1 were not shared by the other short-lived accessions, suggesting that different short-lived seeds express their own sets of genes. These results reinforce the idea that both long-, but also short-lived accessions accumulate sets of transcripts that could enhance longevity.

### Alternative splicing contributes to transcriptome diversity in dry seeds from different accessions

Alternative mRNA splicing (AS) is a mechanism for expanding proteomic diversity and regulating gene expression (Eckardt, 2013), but the contribution of AS events to the landscape of dry seed transcriptomes has not been studied in detail. Given the differences between accessions in the expression of genes belonging to categories related to RNA processing, we performed a systematic analysis of differential AS events using IsoformSwitchAnalyzeR, a recently developed software that enables identification and analysis of alternative splicing and isoform switches with predicted functional consequences (Vitting-Seerup & Sandelin, 2019). The results showed a non-negligible fraction of differential AS events in the dry seed transcriptome between accessions (Figure 5), adding a new level of diversity beyond the expression analysis based on gene levels, and indicating that the differences in stored mRNAs between wild type accessions could have been underestimated. We next predicted the functional consequences of these AS events in every accession compared with Col-0. As observed in Supplemental Figure 2, many genes present important differences with Col-0, including ORF or domain gains, or intron retention differences, reinforcing the idea of a distinct population of alternative-spliced mRNAs among accessions. However, a clear feature associated to longer-lived accessions could not be found.

**Figure 5.**
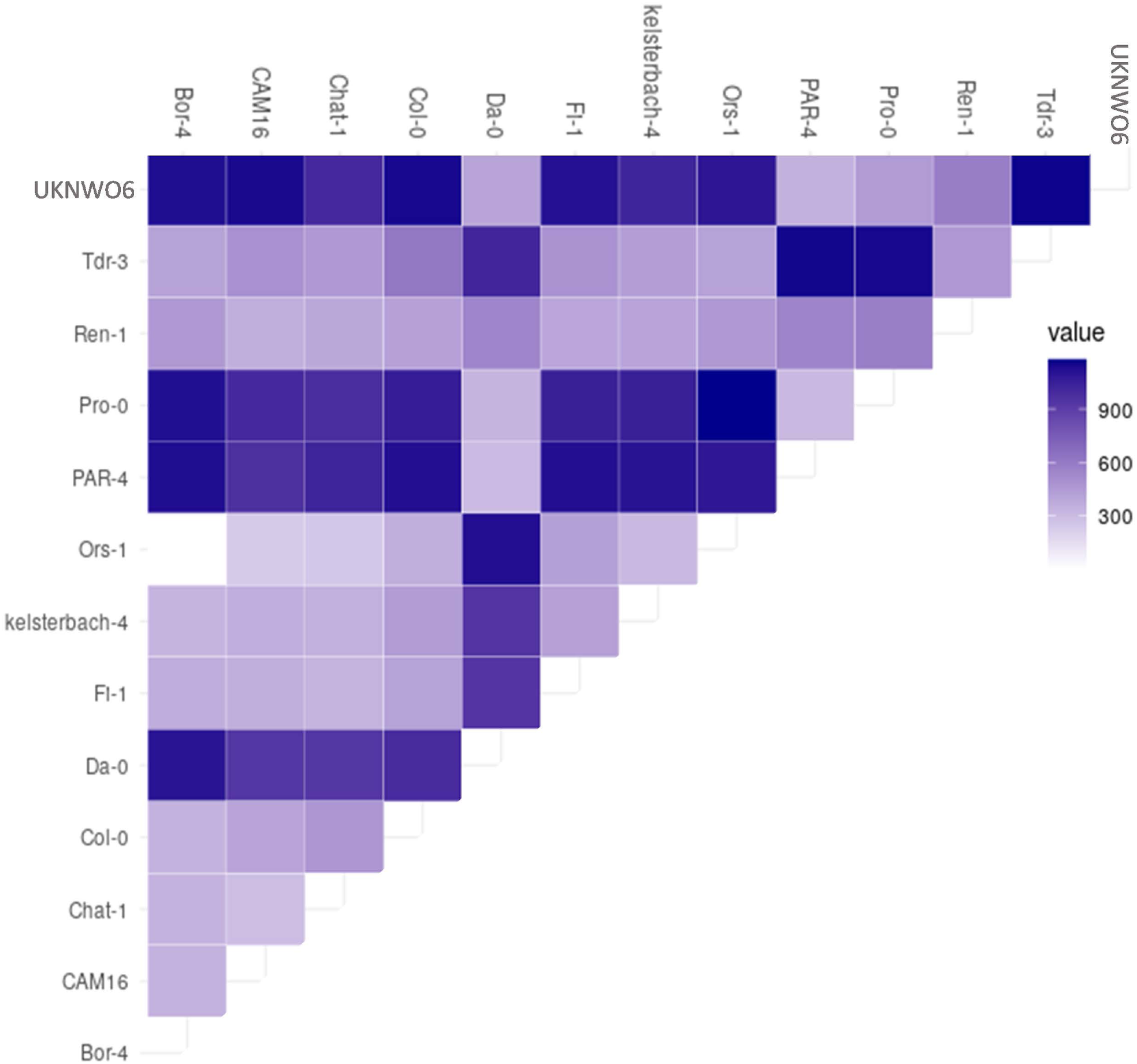
Alternative splicing analysis of the transcriptomes of dry seeds of wild type accessions. Heatmap showing the number of alternative splicing events between accessions according to IsoformSwitchAnalyzeR.

### Reverse genetic analysis identified novel genes involved in seed longevity

To study in more detail the involvement of proteins affecting mRNA regulation in seed longevity, reverse genetics experiments were performed using loss-of-function Arabidopsis lines. Tandem zinc finger protein 9 (TZF9) is an RNA-binding protein involved in the assembly of RNA granules, which harbour transcripts temporally excluded from the translationally active pool. Consequently, this protein influences the number of mRNAs associated with ribosomes (Chantarachot and Bailey-Serres, 2018; Standart and Weil, 2018). TZF9 interacts with poly(A)-binding protein 2 (PAB2), a hallmark constituent of RNA granules (Tabassum et al, 2020), and was more abundant in the Da-0 long-lived accession. Given the relevance of untranslated stored mRNAs in dry seeds, we checked the effect of seed ageing treatments on seeds of the tzf9 mutant. Seeds of tzf9 resisted the CDT better than wild-type seeds. After 21 days of CDT (40°C, 75% RH), 22% of tzf9 seeds, but only 2.6% of wild type seeds, were able to germinate (Figure 6). This highlights the relevance of mRNA packing regulation to seed longevity. Similarly, we tested the response to CDT of mutants in three subunits of the MOS-associated complex (MAC) involved in pre-mRNA splicing (Jia et al, 2017), MOS4, MAC3A and MAC3B. Again, seeds of mos4-1 and the double mutant mac3a/mac3b were able to resist deterioration better than wild type seeds. Whereas 30% of mac3a/mac3b and 18.6% of mos4-1 seeds were able to germinate after 21 days of CDT, only 2.6% of wild types did (Figure 6), uncovering a new role for RNA splicing in shortening seed longevity.

**Figure 6.**
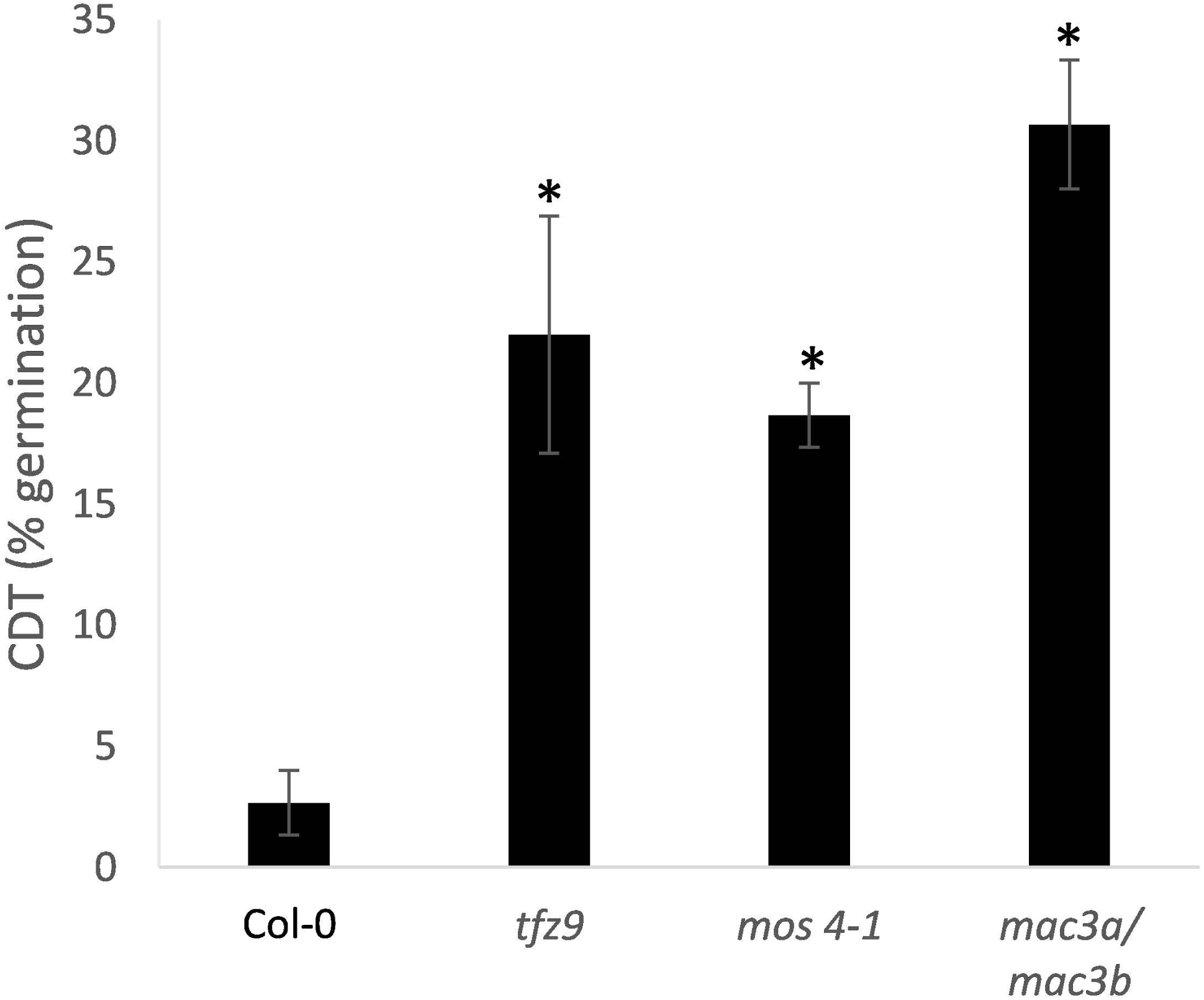
Controlled-deterioration treatment on RNA regulation mutant lines. Percentage of germination at 10 days after sowing of wild-type and tzf9, mac3a/mac3b and mos4-1 seeds aged for 21 days at 75% R.H and 40°C. *Significantly differing from control (Col-0) at P<0.05 (Student’s t-test).

Stored mRNAs of many HSPs particularly accumulated in dry seeds of the long-lived accessions Da-0, Pro-0 and TDr-3 (Supplemental Table 4). RT-PCR data for the highly permeable accession Bs-2 showed that at least HSP70 and HSP17.6A are also more abundant in dry seeds of this accession than in the short-lived Ors-1 (Supplemental Figure 3), indicating that this could be a shared mechanism supporting longevity in highly-permeable seeds. Some of these HSPs are under the control of the heat-shock factors HSF1A/HSF1B (Lohmann et al, 2004; Busch et al, 2004), two transcription factors acting synergistically in the heat-shock response (Li et al, 2010), which accumulate during seed maturation in Arabidopsis (Kotak et al, 2017). Impairment of the HSFA9 expression programme in transgenic tobacco leads to reduced seed longevity (Prieto-Dapena et al, 2006; Tejedor-Cano et al, 2010), but the involvement of HSF1A/HSF1B, and the possible redundancy with other HSFs is unknown. We found that HSP101, HSP70, HSP17.6.A and HSP17.6.C expression in dry seeds was reduced (for HSP70), or practically abolished (for HSP101, HSP17.6.A and HSP17.6.C) in dry seeds of hsf1a/hsf1b, (Figure 7a). These results indicate an absence of redundancy between HSFs for the expression of these particular HSPs during seed maturation that are dependent on HSF1A/HSF1B for expression. Moreover, expression of HSF9A was reduced in hsf1a/hsf1b, indicating that these two transcription factors partially control the HSFA9 expression programme. The CDT (14 d, 40 °C, 75% RH) on hsf1a/hsf1b seeds revealed a clear sensitivity to ageing in the mutant: After 14 days of CDT, hsf1/hsf3 dropped to 37%, whereas 70% of wild-type seeds germinated (Figure 7b), confirming the importance of HSF1A/HSF1B on HSP provision in dry seeds to counteract seed deterioration.

**Figure 7.**
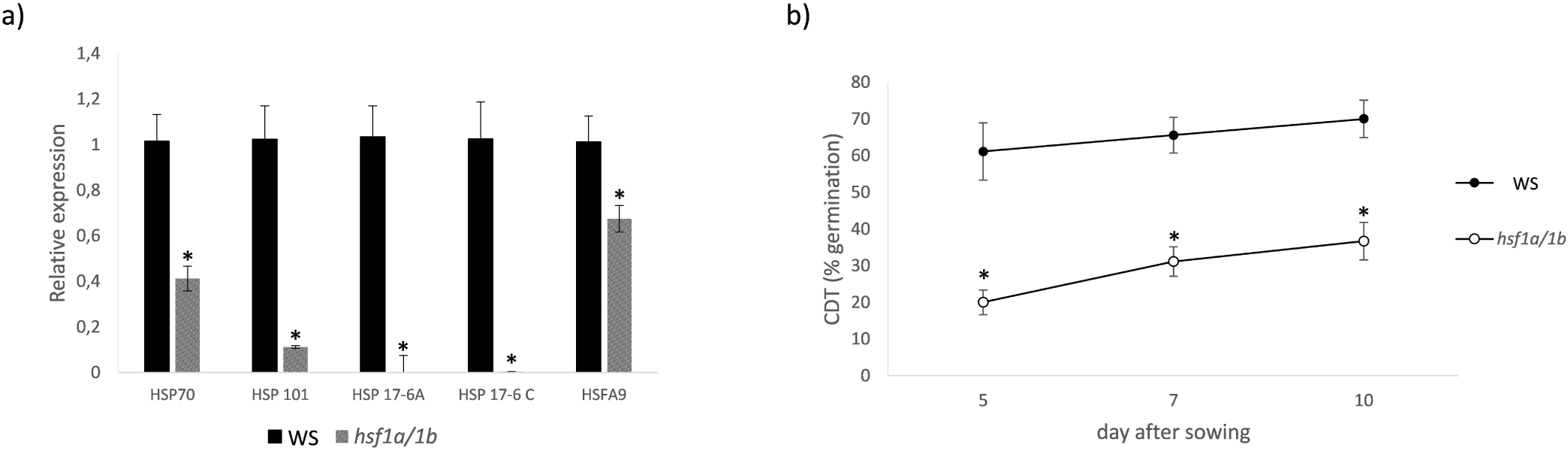
HSPs expression levels and controlled-deterioration treatment of seeds of the hsf1a/hsf1b mutant line. a) Real-time quantitative PCR of HSP70, HSP101, HSP17-6.A and HSP 17-6.C from seeds of the hsf1a/hsf1b Arabidopsis mutant line. Data are the mean of three replicates. Expression for every HSP in wild-type is set to 1 in the y-axis and relative expression in hsf1a/hsf1b is shown. b) Percentage of germination at 5, 7 and 10 days after sowing of wild-type and hsf1a/hsf1b seeds aged for 14 days at 75% R.H and 40°C. * Significantly differing from control (WS) at P<0.05 (Student’s t-test).

### Glutathione concentrations in dry seeds correlated with longevity

The functional category “glutathione metabolic process” was highlighted in clusters 1 and 2 (Figure 4a), which contained genes more expressed in the longer-lived accession Da-0. To investigate the role of this antioxidant in more detail, glutathione concentrations were measured. The accessions investigated here, selected for differences in seed longevity, had clear differences in GSH concentrations and E_GSSG/2GSH_ values before ageing, and these differences correlated with longevity. For example, long-lived accessions (with >40% TG after 30 months ambient ageing or >25% after the CDT (Figure 1) had >0.3 GSH µmol g^-1^ DW (Figure 8a) and E_GSSG_/_2GSH_ values more negative than -188 mV (Figure 8b), whereas shorter-lived accessions (with <5% TG after 30 months ambient ageing or <10% after controlled deterioration) had <0.3 GSH µmol g^-1^ DW and E_GSSG/2GSH_ values more positive than -188 mV. Correlating TG after ambient storage with either pre-storage GSH concentrations, or E_GSSG/2GSH_ values, in all accessions, resulted in Pearson R correlations of 0.79 and -0.78 (Figure 8c and d), respectively, and a trend could also be found when considering TG after CDT of (R=0.64 and []0.80, respectively) (Supplemental Figure 4 a,b). In summary, across a range of Arabidopsis wild-type accessions, concentrations of GSH and E_GSSG/2GSH_ values before ageing provided a predictive indicator of seed longevity.

**Figure 8:**
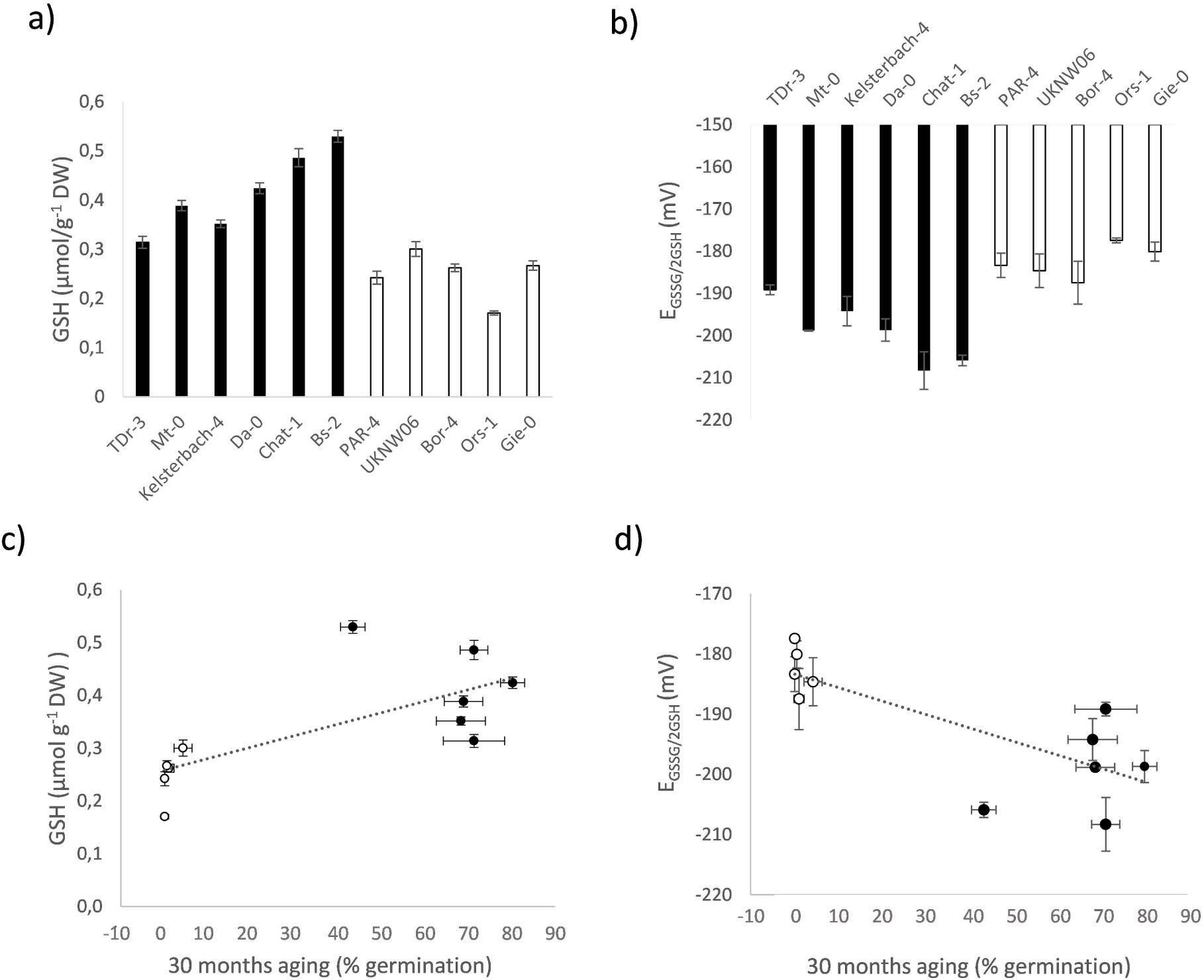
Relationship between glutathione and seed longevity in Arabidopsis wild type accessions. a) Concentrations of glutathione (GSH), and b) glutathione half-cell reduction potentials (E_GSSG_/_2GSH_), reflecting cellular redox state in dry seeds. Data are means of four replicates ± SD. c) and d) Correlation of data shown in a) and b), respectively, with total germination after 30 months of ambient ageing.

## Discussion

Natural conditions that slow down the process of seed ageing, such as deeper soil or cold environments, generate less selective pressure on resistance to ageing than those that accelerate this process, such as shallow burial and warmer climates (Mondoni et al, 2011; Saatkamp et al, 2011; Pierre et al, 2015). Thus, variations in seed longevity of widely dispersed species, such as Arabidopsis thaliana, may reflect adaptive differences to localised habitats. In addition, longevity coexists with related traits, such as dormancy or germination speed, and a delicate balance between them is required to assure the most adequate timing for germination, without penalty to cellular components affecting seed competitiveness (Shen et al, 2011). Adaptation starts the fixation of an advantageous allele for a specific trait, but entails a secondary phase of changes suppressing the pleiotropic effects caused by the first, sometimes detrimental to another trait. Differential gene expression is a mechanism used to compensate these pleiotropic effects (Maisnier-Patin and Andersson, 2004, Pavlicev and Wagner, 2012). When seed properties in natural variations were assessed, positive and negative aspects for seed longevity were found both in long- and short-lived accessions, as one might intuitively expect, as seeds are complex structures with multiple components affecting different traits.

Prevention of germination by dormancy can be alleviated by oxidants, such as H_2_O_2_, and by initial ageing associated with an oxidative shift in the thiol-based cellular redox state (Morscher et al., 2015). Oxidation of seed macromolecules, including nucleotides (mRNA) and to redox-active groups of proteins, may be involved in promoting germination (El-Maarouf-Bouteau et al., 2017; Sano et al., 2020). In fact, T-DNA mutants affected in flavonoid or cutin biosynthesis have more permeable seed coats and are less dormant and less tolerant to ageing (Debeaujon et al. 2000, de Giorgi et al, 2015). By enabling O_2_ diffusion, permeability of the seed coat could be beneficial for germination success if low dormancy suits the local conditions, but would also translate into low longevity. However, an inverse correlation between dormancy and longevity has been observed in various Arabidopsis mutants and natural accessions (Nguyen et al 2012, He et al, 2014). Some of the long-lived accessions studied here, such as Da-0, TDr-3 or Bs-2, showed a much higher permeability than Col-0 (or even higher than some of the shorter-lived accessions, such as PAR-4 or CAM-16, see Figure 2a). This indicates that the detrimental effects on longevity of these permeable seed coats are compensated by other components of the seeds. We found a great number of categories of stored mRNAs enriched upon transcripts more abundant in long-lived accessions. The higher abundance of HSPs in Da-0 and other highly permeable long-lived accessions, such as TDr-3 or Bs-2, could clearly be one of the compensating adaptive changes to this high permeability, and would explain its unexpected longevity. The relevance of these proteins on longevity will be discussed below. Similarly, the high accumulation of transcripts of the catalase CAT2 in Da-0 and TDr-3, or the proline biosynthesis pyrroline-5-carboxylate (P5C) reductase, could also alleviate the higher oxidative stress of highly permeable seeds, at least when seeds were sufficiently hydrated. Catalase could be necessary following seed imbibition, when the reactivation of metabolism generates ROS (Gallardo et al., 2001; Bailly et al.,2002). Moreover, catalase inhibition reduced seed repair after ageing in primed sunflower seeds (Kibinza et al, 2011), highlighting the importance of antioxidant defence in the repair of the damage incurred during ageing. Proline content and P5C reductase levels increase during storage of oat seeds at 28% RH, and has been suggested to play a main role in adaptation to oxidative stress in seeds at high humidity (Kong et al, 2015). Moreover, a high accumulation of transcripts of translational initiation factors in Da-0 could also assure a higher provision of these proteins in the seed, which will guarantee translation efficacy at the time of germination (Dirk and Downie, 2018).

Positive components for longevity were also observed in short-lived seeds, for instance by the accumulation of Lea proteins or ABI5 in the low longevity accessions Ors-1 and Bor-4 (Supplemental Table 3). Lea proteins have been associated with increased seed longevity (Hundertmark et al, 2011), although the molecular mechanism has not been characterised, and abi5 mutant Medicago truncatula plants are sensitive to CDT (Zinsmeister et al, 2016). These observations could in principle be inconsistent, as these accessions are catalogued by us as having low resistance to deterioration, but accumulation of Lea proteins could restrain an otherwise harsher phenotype after ageing in nature. Positive components for longevity in natural accessions presenting short-lived phenotypes were also reported by Sugliani et al, 2009 using introgression lines from the Sei-0 and Sha accessions of Arabidopsis in the abi3-5 and lec1-3 backgrounds, highly impaired in seed longevity. These two accessions are sensitive to deterioration (Renard et al, 2020b), but they could partially re-establish the seed developmental programs controlled by LEC1 and restore the accumulation of seed storage proteins and Lea proteins that were reduced in abi3-5 and lec1-3.

Overall, the differences found in mRNA provision between accessions, both at the level of genes and at the level of spliced variants, could, together with the differences in protein and metabolite provisions (Kliebenstein et al, 2001; Chibani et al, 2006, Joshi et al 2012, Knoch et al, 2017), provide the seed with efficient mechanisms to tune not only its tolerance to deterioration, but other aspects of seed performance, adapting its potential to its particular requirements in nature. This seems reasonable, as the observed natural variability in seed properties such as seed dormancy, germination speed or seed longevity is very high (Alonso-Blanco et al, 2003, Atwell et al, 2010, Renard et al, 2020). Our analysis of stored mRNAs and alternative splicing switches in thirteen different wild type accessions grown under identical environmental conditions clearly indicate that the diversity between accessions is wider than previously assumed (Kimura and Nambara, 2010). From 1600 to 3100 transcripts, and from 360 to 1120 alternative splicing isoforms, depending on the accession, were differentially accumulated with Col-0, suggesting major differences in seed biochemical potential between them.

Alternative splicing can affect the amplitude of expression, the stability and translational efficiency of the mRNA, or it can result in protein isoforms. Alternative transcription start site leads to great differences in translation activity (Rojas-Duran et al, 2012; Ebina et al, 2015; Kurihara et al, 2018). Similarly, alternative termination sites do not seem to increase proteome complexity, but are involved in post-transcriptional regulation (Reyes et al, 2018). The mechanisms by which dry seeds coordinate stored mRNAs translation stalling and protection with priming for efficient translation during imbibition could be mediated by a fine regulation of alternative splicing and by other aspects of RNA regulation, given the differences observed in alternative splicing switches and in expression of genes involved in RNA regulation, but the data using wild type accessions did not give a clear view of the impact of these processes. However, the reverse genetics assays uncover novel roles in longevity for specific genes. Half-life of mRNAs ranges from minutes to hours, suggesting that protective mechanisms to safeguard stored mRNAs during seed life span should exist. Stress granules (SG) (Davies et al, 2012; Chantarachot and Bailey-Serres, 2018) have recently been suggested as one of these protective structures (Sajeev et al., 2019). We found PAB2, a well-stablished marker of SGs, more expressed in the long-lived accession Da-0, Pro-0 or TDr-3 than in the short-lived accessions Ors-1 or Bor-4, suggesting higher accumulation of stress granules that could protect mRNAs from degradation. TZF9 interacts with PAB2 in RNA granules and is important for stalling mRNAs from translation, probably by recruitment in SGs (Tabassum et al, 2019). Unexpectedly, we found tzf9 seeds more resistant to ageing. If SG formation was indeed impaired in tzf9 mutants, this would indicate that RNA protection in SGs is less relevant to longevity than expected. By contrast, ribosome-bound mRNAs were shown to increase in tzf9 (Tabassum et al, 2019). Perhaps this also means that a higher proportion of stored mRNAs is bound to ribosomes in dry seeds. Recently, Bai et al, 2020 showed that ∼60% of stored mRNA in dry seeds was associated to monosomes, suggesting a priming mechanism during seed maturation for specific fates during germination, and an alternative protective mechanism mediated by ribosome protection. This would explain the response obtained in the tzf9 mutant. During germination, protein synthesis requires rapid and selective translation from stored mRNAs, so a ribosome-protected and primed population of mRNAs could confer an advantage to tzf9 seeds after ageing. Similarly, we found in some long-lived accessions higher expression of genes belonging to the MAC complex, involved in the splicing machinery (Jia et al, 2017). Only a small proportion of genes are under the control of the MAC complex for splicing regulation (Jia et al, 2017), discarding a global effect of splicing on the whole population of mRNAs that could impact on seed properties. Seeds of both mac3a/mac3b and mos4-1 mutants performed better than wild type seeds after CDT (see Figure 6), which could indicate that a differentially spliced MAC-regulated gene could confer an advantage to the seeds. The observed increased transcription in the group of long-lived seeds of genes whose loss-of-function seems beneficial to longevity could indicate, in the case of MOS4 and MAC3A/MAC3B, that higher expression of these genes could cause alternative splicing switches in downstream genes that would be beneficial for the seeds in aspects of seed quality or germination other than longevity.

The overabundance of HSP transcripts was highlighted in the highly permeable, long-lived accessions Da-0, Pro-0, TDr-3 or Bs-2. HSP expression is part of normal development during late seed maturation, and its presence is associated with desiccation tolerance, by preventing aggregation of proteins in the low-water content environment of the seed (Gallardo et al., 2001; Wang et al., 2014). In addition, HSPs are important during germination. The cotton GhHSP24.7 modulates cytochrome C/C1 production to induce ROS generation, which accelerates endosperm rupture and promotes seed germination (Ma et al 2019). Transgenic tobacco plants ectopically expressing small HSPs were able to germinate in the dark, indicating that HSPs are part of the light-dependent program during germination (Koo et al, 2015). Moreover, HSPs are ageing-responsive proteins whose abundance increases after natural or artificial ageing treatments (Rajjou et al, 2008; Kaur et al, 2015), and are more expressed in long-lived near isogenic Hordeum vulgare lines (Wozny et al, 2018). A better provision of HSPs in the seed could help refold misfolded proteins during seed desiccation and to foster germination in long-lived aged seeds. In Helianthus annuus, seeds overexpressing the heat-shock factor HSFA9, which over-accumulate HSP101 and small HSPs, showed increased resistance to controlled CDT (Prieto-Dapena et al, 2006), and loss-of-function of HSF9A in tobacco was associated with reduced resistance to artificial ageing (Tejedor-Cano et al, 2010). We show a similar response in the hsf1a/hsf1b double mutant line. Arabidopsis has four members of the HSF1A subfamily, and three of them (HSF1A, 1B and 1E) are expressed during seed development (Kotak et al, 2007), setting up a not well understood multistep mechanism for HSP regulation in the dry seed, similar to the one deployed in heat response (Schöffl et al., 1998) and raising the question of genetic redundancy. The fact that the levels of HSP70, HSP101, HSP 17.6A and 6C dropped dramatically in hsf1a/hsf1b seeds indicates that HSF1A or HSF1B expression is essential for activation of these HSPs. HSP101 is induced by HSF9A in Arabidopsis (Kotak et al, 2007), and repressed in a loss-of-function mutant of HSF9A in tobacco (Tejedor-Cano et al, 2010). HSFA9 expression was reduced in hsf1a/hsf1b seeds (Figure 7a), suggesting that HSF9A is acting in seed development downstream of HSF1A/HSF1B regulating HSPs expression. Interestingly, in the early response to heat stress, HSF1A/HSF1B also mediate the activation of HSF4 (Lohmann et al, 2004), which is also expressed during seed development. In all, these results suggest that the regulation of HSP synthesis during seed maturation, mediated by the HSF1A and 1B transcription factors, could be a novel mechanism to tune seed deterioration in nature.

Seed ageing is associated with the oxidation of macromolecules (Bailly 2004), and efficient antioxidant systems are deployed by the seed to prevent excessive oxidation. The GSSG/GSH redox couple is the main cellular redox buffer in seeds and plays a major role in the regulation of the redox environment. Loss of seed viability was correlated with a shift in the E_GSSG_/_2GSH_ towards more positive values (more oxidising), whereby half of the seed population had lost viability when E_GSSG_/_2GSH_ shifted into a zone of E_GSSG_/_2GSH_ between −180 and −160mV in Pisum sativum, Cytisus scoparius and Lathyrus pratensis (Kranner et al., 2006). This was also observed in barley using 26 genotypes (Nagel et al, 2015) and it is confirmed here for Arabidopsis (Supplemental Figure 4e). Moreover, the data obtained for GSH and E_GSSG_/_2GSH_ in non-aged seeds differing in viability indicate that GSH concentration and E_GSSG_/_2GSH_ in the dry seed are correlated with seed viability (Figure 8c,d and Supplemental Figure 4a,b). This suggests that glutathione biosynthesis, and the ability to maintain a minimum reservoir of GSH before seed drying (>0.3 GSH µmol g^-1^ DW) is a major determinant of viability in wild type accessions of Arabidopsis.

In summary, the genetic complexity of seed longevity in wild type accessions, previously evidenced by the multiple loci identified in numerous genome-association studies (Nagel et al, 2015; Zuo et al, 2019; Renard et al, 2020) is complemented by this study, showing how multiple molecular and cellular factors contribute positively, but also negatively, to this trait, determining the adaptive responses of every accession in nature. Moreover, our results support recent discussions about the impact on seed longevity of the regulation of mRNAs that are stored during seed development, but that will be translated years later.

## Acknowledgments

The authors thank the bioinformatic unit of the IBMCP for technical assistance.

## Authorship

R.N, D.P, P.A, C.R, J.R, R.C, J. F, T.R. and J.G. carried out the experiments. J.G. wrote the manuscript with support from R.N. R.S, E.B. and I.K. helped supervise the project. J.G conceived the original idea.

---

Supplemental Figure 1. Heatmap showing clusters of genes differentially expressed in long-lived accessions (Da-0/Pro-0/Tdr-3) compared with short-lived ones (UKNW06/ Bor-4/ Ors-1/CAM-16). Cluster 1ext: cluster 1 obtained in this experiment including genes mainly overexpressed in long-lived accessions.

Supplemental Figure 2. Consequences of Isoform switch between each accession and Col-0, according to IsoformSwitchAnalyzeR. Number of genes presenting each of these consequences is shown by circle size. For each pair of opposite consequences (e.g. protein domain loss vs gain) (y-axis) this plot shows the fraction of switches, affected by either of the opposing consequence, which results in the consequence indicated (e.g. protein domain loss) (x-axis). If this fraction is significantly different from 0.5, it is shown in red and indicates that there is a systematic bias between the accession and Col-0 in isoform switch consequences.

Supplemental Figure 3. Real-time quantitative PCR of HSP70, HSP101, HSP17-6.A and HSP 17-6.C in seeds of the highly-permeable accessions Bs-2 (representing long-lived) and Ors-1 (representing shorter-lived). Data are means of three replicates. Expression in Ors-1 is set to 1 in the y-axis and relative expression with Bs-2 is shown. * Significantly differing from control (WS) at P<0.05 (Student’s t-test).

Supplemental Figure 4. Correlation between seed longevity and glutathione concentration or glutathione redox potential. Correlation curves of GSH (a) and E_GSSG_/_2GSH_ values (b) of non-aged dry seeds with germination after the controlled deterioration treatment (CDT).

Supplemental Table 2. Samples used for RNAseq and uniquely-mapped reads obtained in each sample after mapping to the TAIR10 Arabidopsis thaliana genome.

Supplemental Table 3. Log_2_ Fold-Change (FC) of gene expression between every accession and Col-0. <u>Diff1</u>: Data is shown in the Excel cell if Log_2_> 0.7 in a long-lived accession. Data is shown in the Excel cell if Log_2_ FC < 0.2 in a short-lived accession. Only genes with Log_2_ FC > 0.7 in all the long-lived accessions with data <u>and</u> Log_2_ FC < 0.2 in all the short-lived accessions with data are listed. <u>Diff4</u>: Data is shown in the Excel cell if Log_2_ FC< -0.7 in a short-lived accession. Data is shown in the Excel cell if Log_2_ FC> -0.2 in a long-lived accession. Only genes with Log_2_ FC< -0.7 in all the long-lived accessions with data <u>and</u> Log_2_ FC > -0.2 in all the short-lived accessions with data are listed.

Supplemental Table 4. Reads-per-kilobase-per million reads (RPKM) data of Da-0, Col-0, Ors-1 and Bor-4 accessions of the 1271 most variable genes (standard deviation > 500).

Supplemental Table 5. Gene ontology (GO) analysis (Biological Process) of genes belonging to clusters 1 and 2 (sheet 1) and from 3 to 6 (sheet 2) in the Da-0 vs. Ors-1/bor-4 comparison, according to Figure 4a. Gene ontology analysis (Biological Process) of genes belonging to clusters 1ext, according to Supplemental Figure 1, in the Da-0/Pro-0/Tdr-3 vs. UKNW06/Ors-1/Bor-4/CAM-16 comparison. GO-acc: GO code; p-value: adjusted p-value after Agrigo v2; FDR: false discovery rate.

**Supplemental Table 1.**
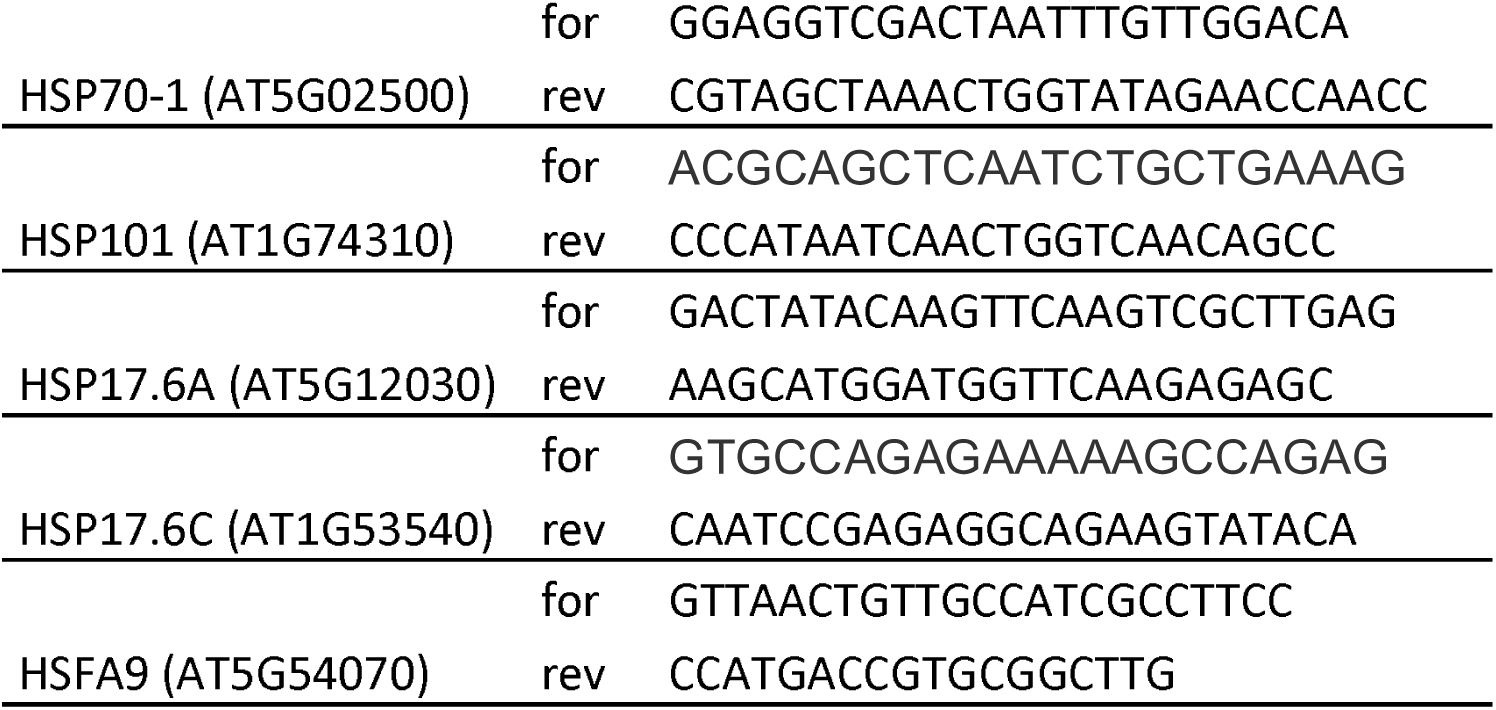
. Primers used for qRT-PCR

## REFERENCES

1. Agacka-Modoch M., Nagel M., Doroszewska T., Lewis R. S., Börner A. Mapping quantitative trait loci determining seed longevity in tobacco (Nicotiana tabacum L.). Euphytica 2015; 202 479–486. 10.1007/s10681-015-1355-x

2. Alonso-Blanco C, Bentsink L, Hanhart CJ, Blankenstijn-de Vries H, Koornneef M Analysis of natural allelic variation at seed dormancy loci of Arabidopsis thaliana. Genetics 2003; 164:711–729.

3. Andrews, S. (2010). FastQC: A Quality Control Tool for High Throughput Sequence Data [Online].Available online at: http://www.bioinformatics.babraham.ac.uk/projects/fastqc/

4. Anders S, Pyl PT, Huber W. HTSeq--a Python framework to work with high-throughput sequencing data. Bioinformatics. 2015 Jan 15;31(2):166–9. doi: 10.1093/bioinformatics/btu638.

5. Angelovici R, Galili G, Fernie AR, Fait A. Seed desiccation: a bridge between maturation and germination. Trends Plant Sci. 2010 Apr;15(4):211–8. doi: 10.1016/j.tplants.2010.01.003. PMID: 20138563.

6. Atwell, S., Huang, Y., Vilhjálmsson, B. et al. Genome-wide association study of 107 phenotypes in Arabidopsis thaliana inbred lines. Nature 2010; 465, 627–631. https://doi.org/10.1038/nature08800

7. Bai, B., van der Horst, S., Cordewener, J., America, T., Hanson, J., & Bentsink, L. Seed-Stored mRNAs that Are Specifically Associated to Monosomes Are Translationally Regulated during Germination. Plant physiology, 2020; 182(1), 378–392. https://doi.org/10.1104/pp.19.00644

8. Bailly, C. Active oxygen species and antioxidants in seed biology 2004. Seed Sci. Res. 14: 93–107.

9. Bailly C, Audigier C, Ladonne F, Wagner MH, Coste F, Corbineau F, Côme D. Changes in oligosaccharide content and antioxidant enzyme activities in developing bean seeds as related to acquisition of drying tolerance and seed quality. J Exp Bot. 2001 Apr;52(357):701–8. doi: 10.1093/jexbot/52.357.701

10. Bissoli G, Niñoles R, Fresquet S, Palombieri S, Bueso E, Rubio L, García-Sánchez MJ, Fernández JA, Mulet JM, Serrano R. Peptidyl-prolyl cis-trans isomerase ROF2 modulates intracellular pH homeostasis in Arabidopsis. Plant J. 2012 May;70(4):704–16. doi: 10.1111/j.1365-313X.2012.04921.x.

11. Bueso E, Ibañez C, Sayas E, Muñoz-Bertomeu J, Gonzalez-Guzmán M, Rodriguez PL, Serrano R. A forward genetic approach in Arabidopsis thaliana identifies a RING-type ubiquitin ligase as a novel determinant of seed longevity. Plant Sci. 2014 Feb;215–216:110-6. doi: 10.1016/j.plantsci.2013.11.004

12. Bueso E, Muñoz-Bertomeu J, Campos F, Brunaud V, Martínez L, Sayas E, Ballester P, Yenush L, Serrano R. ARABIDOPSIS THALIANA HOMEOBOX25 uncovers a role for Gibberellins in seed longevity. Plant Physiol. 2014 Feb;164(2):999–1010. doi: 10.1104/pp.113.232223.

13. Bueso E, Muñoz-Bertomeu J, Campos F, Martínez C, Tello C, Martínez-Almonacid I, Ballester P, Simón-Moya M, Brunaud V, Yenush L, Ferrándiz C, Serrano R. Arabidopsis COGWHEEL1 links light perception and gibberellins with seed tolerance to deterioration. Plant J. 2016 Sep;87(6):583–96. doi: 10.1111/tpj.13220.

14. Buitink, J. and Leprince, O. Intracellular glasses and seed survival in the dry state. C. R. Biol. . 2008; 331: 788–95.

15. Busch W, Wunderlich M, Schöffl F. Identification of novel heat shock factor-dependent genes and biochemical pathways in Arabidopsis thaliana. Plant J. 2005 Jan;41(1):1–14. doi: 10.1111/j.1365-313X.2004.02272.x.

16. Chantarachot T, Bailey-Serres J. Polysomes, Stress Granules, and Processing Bodies: A Dynamic Triumvirate Controlling Cytoplasmic mRNA Fate and Function. Plant Physiol. 2018 Jan;176(1):254–269. doi: 10.1104/pp.17.01468.

17. Chen X, Yin G, Börner A, Xin X, He J, Nagel M, Liu X, Lu X. Comparative physiology and proteomics of two wheat genotypes differing in seed storage tolerance. Plant Physiol Biochem. 2018 Sep;130:455–463. doi: 10.1016/j.plaphy.2018.07.022

18. Chibani, K., Ali-Rachedi, S., Job, C., Job, D., Jullien, M., & Grappin, P. (2006). Proteomic analysis of seed dormancy in Arabidopsis. Plant physiology, 142(4), 1493–1510. https://doi.org/10.1104/pp.106.087452

19. Davies E, Stankovic B, Vian A, Wood AJ. Where has all the message gone? Plant Sci. 2012 Apr;185–186:23-32. doi: 10.1016/j.plantsci.2011.08.001.

20. Debeaujon, I., Leon-Kloosterziel, K.M. and Koornneef, M. (2000) Influence of the testa on seed dormancy, germination, and longevity in Arabidopsis. Plant Physiol. 122: 403– 414.

21. De Giorgi J, Piskurewicz U, Loubery S, Utz-Pugin A, Bailly C, Mène-Saffrané L, Lopez-Molina L. An Endosperm-Associated Cuticle Is Required for Arabidopsis Seed Viability, Dormancy and Early Control of Germination. PLoS Genet. 2015 Dec 17;11(12):e1005708. doi: 10.1371/journal.pgen.1005708

22. Demonsais L, Utz-Pugin A, Loubéry S, Lopez-Molina L. Identification of tannic cell walls at the outer surface of the endosperm upon Arabidopsis seed coat rupture. Plant J. 2020 Nov;104(3):567–580. doi: 10.1111/tpj.14994.

23. Dirk LMA, Downie AB). An examination of Job’s rule: protection and repair of the proteins of the translational apparatus in seeds. Seed Science Research 2018; 8,168– 181. https://doi.org/10.1017/S0960258518000284

24. Ebina I, Takemoto-Tsutsumi M, Watanabe S, Koyama H, Endo Y, Kimata K, Igarashi T, Murakami K, Kudo R, Ohsumi A, Noh AL, Takahashi H, Naito S, Onouchi H. Identification of novel Arabidopsis thaliana upstream open reading frames that control expression of the main coding sequences in a peptide sequence-dependent manner. Nucleic Acids Res. 2015 Feb 18;43(3):1562–76. doi: 10.1093/nar/gkv018.

25. El-Maarouf-Bouteau H, Meimoun P, Job C, Job D, Bailly C. Role of protein and mRNA oxidation in seed dormancy and germination. Front Plant Sci. 2013 Apr 8;4:77. doi: 10.3389/fpls.2013.00077.

26. Gallardo K, Job C, Groot SP, Puype M, Demol H, Vandekerckhove J, Job D. Proteomic analysis of arabidopsis seed germination and priming. Plant Physiol. 2001 Jun;126(2):835–48. doi: 10.1104/pp.126.2.835.

27. He H, de Souza Vidigal D, Snoek LB, Schnabel S, Nijveen H, Hilhorst H, Bentsink L. Interaction between parental environment and genotype affects plant and seed performance in Arabidopsis. J Exp Bot. 2014 Dec;65(22):6603–15. doi:10.1093/jxb/eru378.

28. Hundertmark M. Buitink J. Leprince O. Hincha D.K. The reduction of seed-specific dehydrins reduces seed longevity in Arabidopsis thaliana. Seed Sci. Res. 2011; 21 : 165–173 .

29. Jia T, Zhang B, You C, Zhang Y, Zeng L, Li S, Johnson KCM, Yu B, Li X, Chen X. The Arabidopsis MOS4-Associated Complex Promotes MicroRNA Biogenesis and Precursor Messenger RNA Splicing. Plant Cell. 2017 Oct;29(10):2626–2643. doi: 10.1105/tpc.17.00370.

30. Joshi HJ, Christiansen KM, Fitz J, Cao J, Lipzen A, Martin J, Smith-Moritz AM, Pennacchio LA, Schackwitz WS, Weigel D, Heazlewood JL. 1001 Proteomes: a functional proteomics portal for the analysis of Arabidopsis thaliana accessions. Bioinformatics. 2012 May 15;28(10):1303-6. doi: 10.1093/bioinformatics/bts133

31. Kaur H, Petla BP, Kamble NU, Singh A, Rao V, Salvi P, Ghosh S, Majee M. Differentially expressed seed ageing responsive heat shock protein OsHSP18.2 implicates in seed vigor, longevity and improves germination and seedling establishment under abiotic stress. Front Plant Sci. 2015 Sep 14;6:713. doi: 10.3389/fpls.2015.00713

32. Kibinza S, Bazin J, Bailly C, Farrant J, Corbineau F, El-Maarouf-Bouteau H, Catalase is a key enzyme in seed recovery from ageing during priming, Plant Science, Volume 181, Issue 3, 2011, Pages 309-315, ISSN 0168-9452,https://doi.org/10.1016/j.plantsci.2011.06.003.

33. Kim D, Langmead B and Salzberg SL. HISAT: a fast spliced aligner with low memory requirements. Nat Methods 12, 357–360 (2015). https://doi.org/10.1038/nmeth.3317

34. Kimura M, Nambara E. Stored and neosynthesized mRNA in Arabidopsis seeds: effects of cycloheximide and controlled deterioration treatment on the resumption of transcription during imbibition. Plant Mol Biol. 2010 May;73(1-2):119–29. doi: 10.1007/s11103-010-9603-x.

35. Kliebenstein DJ, Kroymann J, Brown P, Figuth A, Pedersen D, Gershenzon J, Mitchell-Olds T. Genetic control of natural variation in Arabidopsis glucosinolate accumulation. Plant Physiol. 2001 Jun;126(2):811–25. doi: 10.1104/pp.126.2.811.

36. Knoch, D., Riewe, D., Meyer, R. C., Boudichevskaia, A., Schmidt, R., & Altmann, T. (2017). Genetic dissection of metabolite variation in Arabidopsis seeds: evidence for mQTL hotspots and a master regulatory locus of seed metabolism. Journal of experimental botany, 68(7), 1655–1667. https://doi.org/10.1093/jxb/erx049

37. Kong, L., Huo, H., & Mao, P. Antioxidant response and related gene expression in aged oat seed. Frontiers in plant science, 2015; 6, 158. https://doi.org/10.3389/fpls.2015.00158

38. Koo H. J, Park S. M, Kim K. P, Suh M. C, Lee M. O, Lee S, Xinli X, Hong C. B. Small Heat Shock Proteins Can Release Light Dependence of Tobacco Seed during Germination, Plant Physiology Mar 2015, 167 (3) 1030–1038; DOI: 10.1104/pp.114.252841

39. Kotak S., Vierling E., Baumlein H. & Koskull-Döring P. (2007) A novel transcriptional cascade regulating expression of heat stress proteins during seed development of Arabidopsis. The Plant Cell 19, 182–195.

40. Kranner I. 1998. Determination of Glutathione, Glutathione Disulphide and Two Related Enzymes,Glutathione Reductase and Glucose-6-Phosphate Dehydrogenase, in Fungal and Plant Cells. In: Varma A, ed. Springer Lab Manual. Mycorrhiza Manual. Berlin, Heidelberg: Springer, 227–241.z

41. Kranner I, Birtic S. A modulating role for antioxidants in desiccation tolerance. Integr Comp Biol. 2005 Nov;45(5):734–40. doi: 10.1093/icb/45.5.734.

42. Kranner I, Birtić S, Anderson KM, Pritchard HW. Glutathione half-cell reduction potential: a universal stress marker and modulator of programmed cell death? Free Radic Biol Med. 2006 Jun 15;40(12):2155–65. doi: 10.1016/j.freeradbiomed.2006.02.013.

43. Kurihara Y, Makita Y, Kawashima M, Fujita T, Iwasaki S, Matsui M. Transcripts from downstream alternative transcription start sites evade uORF-mediated inhibition of gene expression in Arabidopsis. Proc Natl Acad Sci U S A. 2018 Jul 24;115(30):7831–7836. doi: 10.1073/pnas.1804971115.

44. Kusunoki K, Nakano Y, Tanaka K, Sakata Y, Koyama H, Kobayashi Y. Transcriptomic variation among six Arabidopsis thaliana accessions identified several novel genes controlling aluminium tolerance. Plant Cell Environ. 2017 Feb;40(2):249–263. doi: 10.1111/pce.12866.

45. Leon-Kloosterziel KM, Keijzer CJ, Koornneef M. A Seed Shape Mutant of Arabidopsis That Is Affected in Integument Development. Plant Cell. 1994 Mar;6(3):385–392. doi: 10.1105/tpc.6.3.385. PMID: 12244241; PMCID: PMC160441.

46. Li Z, Wu S, Chen J, Wang X, Gao J, Ren G, Kuai B. NYEs/SGRs-mediated chlorophyll degradation is critical for detoxification during seed maturation in Arabidopsis. Plant J. 2017 Nov;92(4):650–661. doi: 10.1111/tpj.13710.

47. Li M, Berendzen KW, Schöffl F. Promoter specificity and interactions between early and late Arabidopsis heat shock factors. Plant Mol Biol. 2010 Jul;73(4-5):559–67. doi: 10.1007/s11103-010-9643-2.

48. Lohmann, C., Eggers-Schumacher, G., Wunderlich, M. et al. Two different heat shock transcription factors regulate immediate early expression of stress genes in Arabidopsis . Mol Genet Genomics 271, 11–21 (2004). https://doi.org/10.1007/s00438-003-0954-8

49. Love MI, Huber W, Anders S (2014). “Moderated estimation of fold change and dispersion for RNA-seq data with DESeq2.” Genome Biology, 15, 550. doi: 10.1186/s13059-014-0550-8.

50. Ma W, Guan X, Li J, Pan R, Wang L, Liu F, Ma H, Zhu S, Hu J, Ruan YL, Chen X, Zhang T. Mitochondrial small heat shock protein mediates seed germination via thermal sensing. Proc Natl Acad Sci U S A. 2019 Mar 5;116(10):4716–4721. doi: 10.1073/pnas.1815790116

51. Maisnier-Patin S, Andersson DI. Adaptation to the deleterious effects of antimicrobial drug resistance mutations by compensatory evolution. Res Microbiol. 2004 Jun;155(5):360–9. doi: 10.1016/j.resmic.2004.01.019

52. Maldonado-Bonilla LD, Eschen-Lippold L, Gago-Zachert S, Tabassum N, Bauer N, Scheel D, Lee J. The Arabidopsis tandem zinc finger 9 protein binds RNA and mediates pathogen-associated molecular pattern-triggered immune responses. Plant Cell Physiol. 2014 Feb;55(2):412–25. doi: 10.1093/pcp/pct175.

53. Martin, M. Cutadapt removes adapter sequences from high-throughput sequencing reads. EMBnet.journal, [S.l.], v. 17, n. 1, p. pp. 10-12, may 2011. ISSN 2226-6089.

54. Meßner, B., Thulke, O. & Schäffner, A.R. Arabidopsis glucosyltransferases with activities toward both endogenous and xenobiotic substrates. Planta 217, 138–146 (2003). https://doi.org/10.1007/s00425-002-0969-0

55. Molina I, Ohlrogge JB, Pollard M. Deposition and localization of lipid polyester in developing seeds of Brassica napus and Arabidopsis thaliana. Plant J. 2008 Feb;53(3):437–49. doi: 10.1111/j.1365-313X.2007.03348.x.

56. Mondoni A, Probert RJ, Rossi G, Vegini E, Hay FR. Seeds of alpine plants are short lived: implications for long-term conservation. Ann Bot. 2011 Jan;107(1):171–9. doi: 10.1093/aob/mcq222.

57. Monte E, Tepperman JM, Al-Sady B, Kaczorowski KA, Alonso JM, Ecker JR, Li X, Zhang Y, Quail PH. The phytochrome-interacting transcription factor, PIF3, acts early, selectively, and positively in light-induced chloroplast development. Proc Natl Acad Sci U S A. 2004 Nov 16;101(46):16091–8. doi: 10.1073/pnas.0407107101.

58. Morscher F, Kranner I, Arc E, Bailly C, Roach T. Glutathione redox state, tocochromanols, fatty acids, antioxidant enzymes and protein carbonylation in sunflower seed embryos associated with after-ripening and ageing. Ann Bot. 2015 116: 669–678. https://doi.org/10.1093/aob/mcv108

59. Nagel M, Kranner I, Neumann K, Rolletschek H, Seal CE, Colville L, Fernández-Marín B, Börner A. Genome-wide association mapping and biochemical markers reveal that seed ageing and longevity are intricately affected by genetic background and developmental and environmental conditions in barley. Plant Cell Environ. 2015 Jun;38(6):1011–22. doi: 10.1111/pce.12474.

60. Nakabayashi, K., Okamoto, M., Koshiba, T., Kamiya, Y. and Nambara, E. Genome-wide profiling of stored mRNA in Arabidopsis thaliana seed germination: epigenetic and genetic regulation of transcription in seed. Plant J. 2005; 41: 697–709.

61. Nakajima S, Ito H, Tanaka R, Tanaka A. Chlorophyll b reductase plays an essential role in maturation and storability of Arabidopsis seeds. Plant Physiol. 2012 Sep;160(1):261–73. doi: 10.1104/pp.112.196881.

62. Nakashima K, Fujita Y, Kanamori N, Katagiri T, Umezawa T, Kidokoro S, Maruyama K, Yoshida T, Ishiyama K, Kobayashi M, Shinozaki K, Yamaguchi-Shinozaki K. Three Arabidopsis SnRK2 protein kinases, SRK2D/SnRK2.2, SRK2E/SnRK2.6/OST1 and SRK2I/SnRK2.3, involved in ABA signaling are essential for the control of seed development and dormancy. Plant Cell Physiol. 2009 Jul;50(7):1345–63. doi: 10.1093/pcp/pcp083.

63. Nguyen TP, Keizer P, van Eeuwijk F, Smeekens S, Bentsink L. Natural variation for seed longevity and seed dormancy are negatively correlated in Arabidopsis. Plant Physiol. 2012 Dec;160(4):2083–92. doi: 10.1104/pp.112.206649.

64. Nguyen, T.P., Cueff, G., Hegedus, D.D., Rajjou, L. and Bentsink, L. A role for seed storage proteins in Arabidopsis seed longevity. J. Exp. Bot. 2015; 66: 6399–6413. Oñate-Sánchez L, Vicente-Carbajosa J. 2008. DNA-free RNA isolation protocols for Arabidopsis thaliana, including seeds and siliques. BMC Research Notes 1: 93.

65. Patro, R., Duggal, G., Love, M. et al. Salmon provides fast and bias-aware quantification of transcript expression. Nat Methods 14, 417–419 (2017). https://doi.org/10.1038/nmeth.4197

66. Pavlicev M, Wagner GP. A model of developmental evolution: selection, pleiotropy and compensation. Trends Ecol Evol. 2012 Jun;27(6):316–22. doi: 10.1016/j.tree.2012.01.016.

67. Pellizzaro A, Neveu M, Lalanne D, Ly Vu B, Kanno Y, Seo M, Leprince O, Buitink J. A role for auxin signaling in the acquisition of longevity during seed maturation. New Phytol. 2020 Jan;225(1):284–296. doi: 10.1111/nph.16150

68. Pierre JS, Perroux J, Whan A, Rae AL, Bonnett GD. Poor Fertility, Short Longevity, and Low Abundance in the Soil Seed Bank Limit Volunteer Sugarcane from Seed. Front Bioeng Biotechnol. 2015 Jun 4;3:83. doi: 10.3389/fbioe.2015.00083

69. Prieto-Dapena P, Castaño R, Almoguera C, Jordano J. Improved resistance to controlled deterioration in transgenic seeds. Plant Physiol. 2006 Nov;142(3):1102–12. doi: 10.1104/pp.106.087817.

70. Probert R., Adams J., Coneybeer J., Crawford A., Hay F. Seed quality for conservation is critically affected by pre-storage factors. Australian Journal of Botany 2007; 55, 326–335. https://doi.org/10.1071/BT06046

71. Rajjou L, Lovigny Y, Groot SP, Belghazi M, Job C, Job D. Proteome-wide characterization of seed ageing in Arabidopsis: a comparison between artificial and natural ageing protocols. Plant Physiol. 2008 Sep;148(1):620–41. doi: 10.1104/pp.108.123141

72. Renard J, Niñoles R, Martínez-Almonacid I, Gayubas B, Mateos-Fernández R, Bissoli G, Bueso E, Serrano R, Gadea J. Identification of novel seed longevity genes related to oxidative stress and seed coat by genome-wide association studies and reverse genetics. Plant Cell Environ. 2020 Oct;43(10):2523–2539. doi: 10.1111/pce.13822.

73. Renard J, Martínez-Almonacid I, Sonntag A, Molina I, Moya-Cuevas J, Bissoli G, Muñoz-Bertomeu J, Faus I, Niñoles R, Shigeto J, Tsutsumi Y, Gadea J, Serrano R, Bueso E. PRX2 and PRX25, peroxidases regulated by COG1, are involved in seed longevity in Arabidopsis. Plant Cell Environ. 2020 Feb;43(2):315–326. doi: 10.1111/pce.13656.

74. Renard J, Martínez-Almonacid I, Queralta Castillo I, Sonntag A, Hashim A, Bissoli G, Campos L, Muñoz-Bertomeu J, Niñoles R, Roach T, Sánchez-León S, Ozuna CV, Gadea J, Lisón P, Kranner I, Barro F, Serrano R, Molina I, Bueso E. Apoplastic lipid barriers regulated by conserved homeobox transcription factors extend seed longevity in multiple plant species. New Phytol. 2021 Jul;231(2):679–694. doi: 10.1111/nph.17399.

75. Reyes A, Huber W. Alternative start and termination sites of transcription drive most transcript isoform differences across human tissues. Nucleic Acids Res. 2018 Jan 25;46(2):582–592. doi: 10.1093/nar/gkx1165

76. Righetti, K., Ly Vu, J., Pelletier, S., Ly Vu, B., Glaab, E., Lalanne, D., et al. Inference of longevity-related genes from a robust co-expression network of seed maturation identifies new regulators linking seed storability to biotic defense-related pathways. Plant Cell 2015; 27: 2692–2708.

77. Ringli C, Bigler L, Kuhn BM, et al. The modified flavonol glycosylation profile in the Arabidopsis rol1 mutants results in alterations in plant growth and cell shape formation. Plant Cell. 2008;20(6):1470–1481. doi:10.1105/tpc.107.053249

78. Roach T, Beckett RP, Minibayeva FV, Colville L, Whitaker C, Chen H, Bailly C, Kranner I. Extracellular superoxide production, viability and redox poise in response to desiccation in recalcitrant Castanea sativa seeds. Plant Cell Environ. 2010 Jan;33(1):59–75. doi: 10.1111/j.1365-3040.2009.02053.x.

79. Rojas-Duran, M. F., & Gilbert, W. V. Alternative transcription start site selection leads to large differences in translation activity in yeast. RNA (New York, N.Y.) 2012; 18(12), 2299–2305. https://doi.org/10.1261/rna.035865.112

80. Saatkamp, A., Affre, L., Baumberger, T., Dumas, P.-J., Gasmi, A., Gachet, S., et al. Soil depth detection by seeds and diurnally fluctuating temperatures: different dynamics in 10 annual plants. Plant Soil 2011; 349, 331–340. doi:10.1007/s11104-011-0878-8

81. Sajeev N, Bai B, Bentsink L. Seeds: A Unique System to Study Translational Regulation. Trends Plant Sci. 2019 Jun;24(6):487–495. doi: 10.1016/j.tplants.2019.03.011

82. Sano N, Rajjou L, North HM. Lost in Translation: Physiological Roles of Stored mRNAs in Seed Germination. Plants (Basel). 2020 Mar 10;9(3):347. doi: 10.3390/plants9030347.

83. Sano N, Kim JS, Onda Y, Nomura T, Mochida K, Okamoto M, Seo M. RNA-Seq using bulked recombinant inbred line populations uncovers the importance of brassinosteroid for seed longevity after priming treatments. Sci Rep. 2017 Aug 14;7(1):8095. doi: 10.1038/s41598-017-08116-5.

84. Sano N, Rajjou L, North HM, Debeaujon I, Marion-Poll A, Seo M. Staying Alive: Molecular Aspects of Seed Longevity. Plant Cell Physiol. 2016 Apr;57(4):660–74. doi: 10.1093/pcp/pcv186.

85. Sattler SE, Gilliland LU, Magallanes-Lundback M, Pollard M, DellaPenna D. Vitamin E is essential for seed longevity and for preventing lipid peroxidation during germination. Plant Cell. 2004 Jun;16(6):1419–32. doi: 10.1105/tpc.021360.

86. Schöffl F, Prändl R, Reindl A. Regulation of the heat-shock response. Plant Physiol. 1998 Aug;117(4):1135–41. doi: 10.1104/pp.117.4.1135

87. Schramm F, Larkindale J, Kiehlmann E, Ganguli A, Englich G, Vierling E, von Koskull-Döring P. A cascade of transcription factor DREB2A and heat stress transcription factor HsfA3 regulates the heat stress response of Arabidopsis. Plant J. 2008 Jan;53(2):264–74. doi: 10.1111/j.1365-313X.2007.03334.x.

88. Shao Q, Liu X, Su T, Ma C, Wang P. New Insights Into the Role of Seed Oil Body Proteins in Metabolism and Plant Development. Front Plant Sci. 2019;10:1568. doi:10.3389/fpls.2019.01568

89. Shen Y, Zhao C, Liu W, Seed vigor and plant competitiveness resulting from seeds of Eupatorium adenophorum in a persistent soil seed bank, Flora - Morphology, Distribution, Functional Ecology of Plants 2011 Volume 206, Issue 11, , Pages 935–942, ISSN 0367-2530, https://doi.org/10.1016/j.flora.2011.07.002.

90. Schafer,F.Q. and Buettner G.R. Redox environment of the cell as viewed through the redox state of the glutathione disulphide/glutathione couple. Free Radic. Biol. Med., 30 (2001), pp. 1191–1212. doi: 10.1016/s0891-5849(01)00480-4

91. Standart N, Weil D. P-Bodies: Cytosolic Droplets for Coordinated mRNA Storage. Trends Genet. 2018 Aug;34(8):612–626. doi: 10.1016/j.tig.2018.05.005.

92. Sugliani M, Rajjou L, Clerkx EJ, Koornneef M, Soppe WJ. Natural modifiers of seed longevity in the Arabidopsis mutants abscisic acid insensitive3-5 (abi3-5) and leafy cotyledon1-3 (lec1-3). New Phytol. 2009 Dec;184(4):898–908. doi: 10.1111/j.1469-8137.2009.03023.x.

93. Tabassum N, Eschen-Lippold L, Athmer B, Baruah M, Brode M, Maldonado-Bonilla LD, Hoehenwarter W, Hause G, Scheel D, Lee J. Phosphorylation-dependent control of an RNA granule-localized protein that fine-tunes defence gene expression at a post-transcriptional level. Plant J. 2020 Mar;101(5):1023–1039. doi: 10.1111/tpj.14573.

94. Tejedor-Cano J, Prieto-Dapena P, Almoguera C, Carranco R, Hiratsu K, Ohme-Takagi M, Jordano J. Loss of function of the HSFA9 seed longevity program. Plant Cell Environ. 2010 Aug 1;33(8):1408–17. doi: 10.1111/j.1365-3040.2010.02159.x.

95. van Veen H, Vashisht D, Akman M, Girke T, Mustroph A, Reinen E, Hartman S, Kooiker M, van Tienderen P, Schranz ME, Bailey-Serres J, Voesenek LA, Sasidharan R. Transcriptomes of Eight Arabidopsis thaliana Accessions Reveal Core Conserved, Genotype- and Organ-Specific Responses to Flooding Stress. Plant Physiol. 2016 Oct;172(2):668–689. doi: 10.1104/pp.16.00472.

96. Vitting-Seerup K, Sandelin A. IsoformSwitchAnalyzeR: analysis of changes in genome-wide patterns of alternative splicing and its functional consequences. Bioinformatics. 2019 Nov 1;35(21):4469–4471. doi: 10.1093/bioinformatics/btz247.

97. Walters C, Wheeler L, Stanwood PC. Longevity of cryogenically stored seeds. Cryobiology. 2004 Jun;48(3):229–44. doi: 10.1016/j.cryobiol.2004.01.007.15157772.

98. Wang W, He A, Peng S, Huang J, Cui K, Nie L. The Effect of Storage Condition and Duration on the Deterioration of Primed Rice Seeds. Front Plant Sci. 2018; 9:172. doi:10.3389/fpls.2018.00172

99. Wang WQ, Ye JQ, Rogowska-Wrzesinska A, Wojdyla KI, Jensen ON, Møller IM, Song SQ. Proteomic comparison between maturation drying and prematurely imposed drying of Zea mays seeds reveals a potential role of maturation drying in preparing proteins for seed germination, seedling vigor, and pathogen resistance. J Proteome Res. 2014 Feb 7;13(2):606–26. doi: 10.1021/pr4007574

100. Zhang, R., Calixto, C.P.G., Marquez, Y., Venhuizen, P., Tzioutziou, N.A., Guo, W., Spensley, M., Frei dit Frey, N., Hirt, H., James, A.B., Nimmo, H.G., Barta, A., Kalyna, M., Brown, J.W.S (2016) AtRTD2: A Reference Transcript Dataset for accurate quantification of alternative splicing and expression changes in Arabidopsis thaliana RNA-seq data. bioRxiv doi: http://dx.doi.org/10.1101/051938

101. Zinsmeister, J., Lalanne, D., Terrasson, E., Chatelain, E., Vandecasteele, C., Vu, B. L., Dubois-Laurent, C., Geoffriau, E., Signor, C. L., Dalmais, M., Gutbrod, K., Dörmann, P., Gallardo, K., Bendahmane, A., Buitink, J., & Leprince, O. (2016). ABI5 Is a Regulator of Seed Maturation and Longevity in Legumes. The Plant cell, 28(11), 2735–2754. https://doi.org/10.1105/tpc.16.00470

102. Zuo JH, Chen FY, Li XY, Xia XC, Cao H, Liu JD, Liu YX Genome-wide association study reveals loci associated with seed longevity in common wheat (Triticum aestivum L.). Plant Breed, 2019. https://doi.org/10.1111/pbr.12784

