## Supplementary figures and images for "Comparative analysis of wild type accessions reveals novel determinants of Arabidopsis seed longevity"

### Supplemental Figure 1

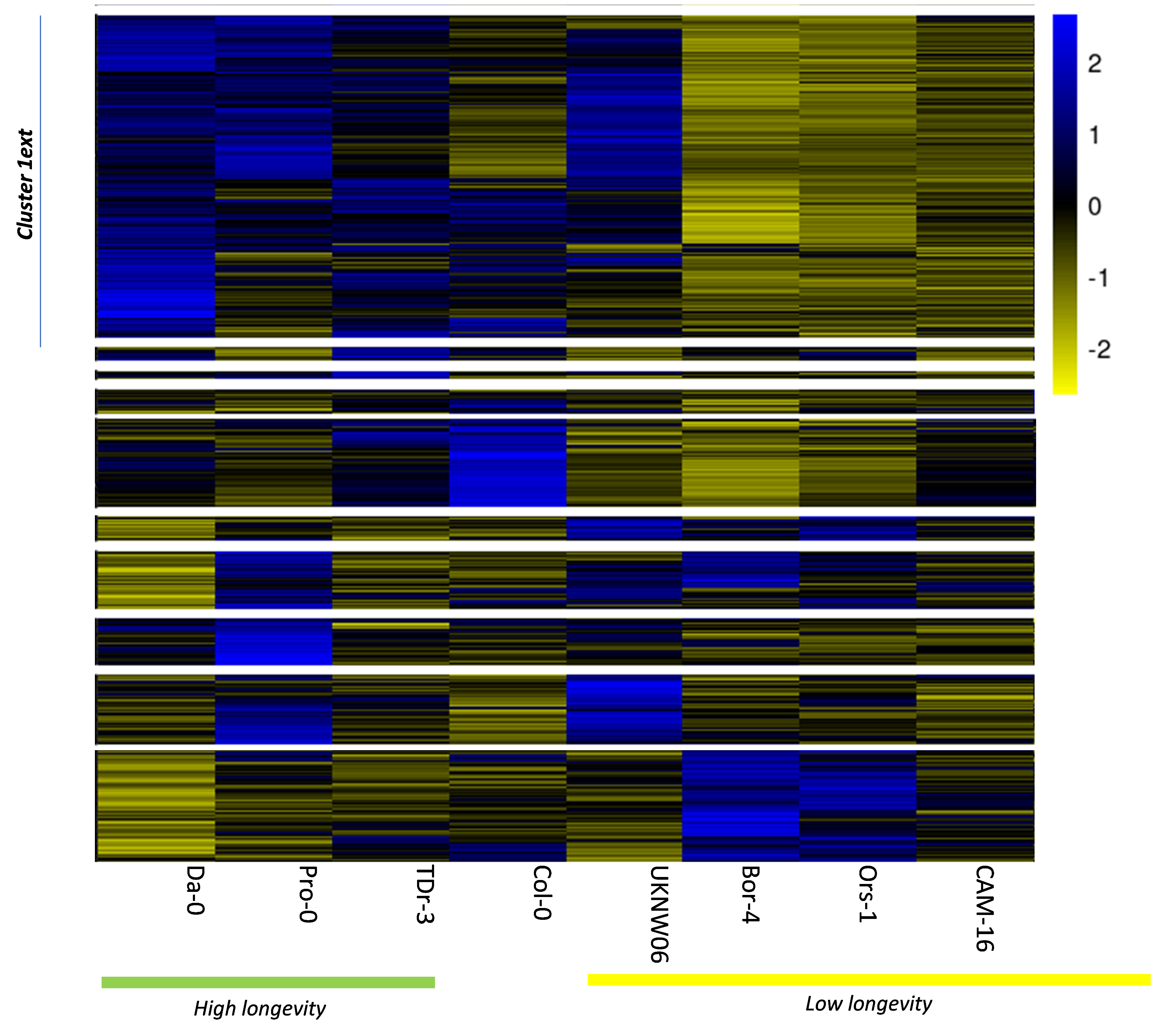

### Supplemental Figure 2

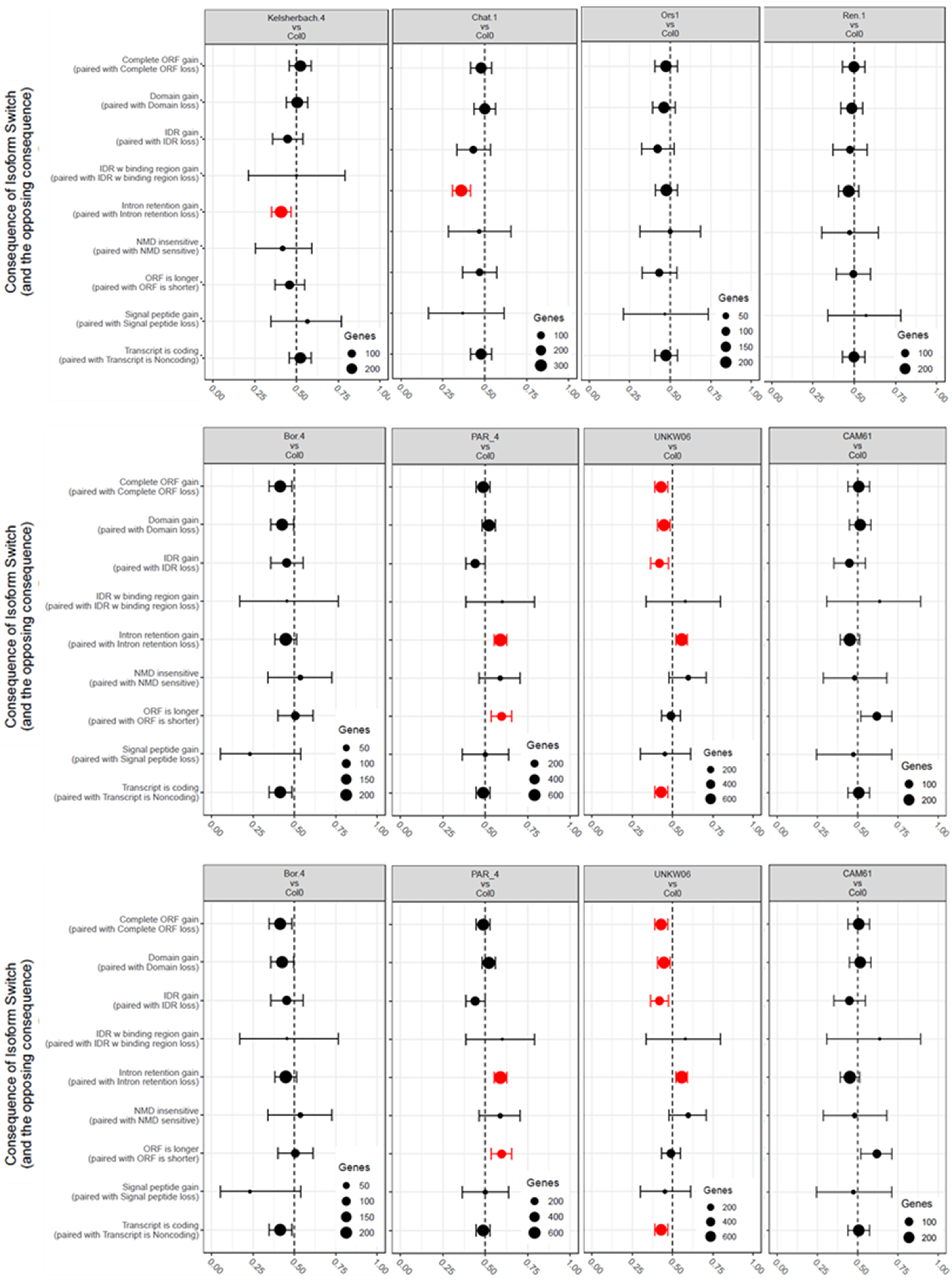

### Supplemental Figure 3

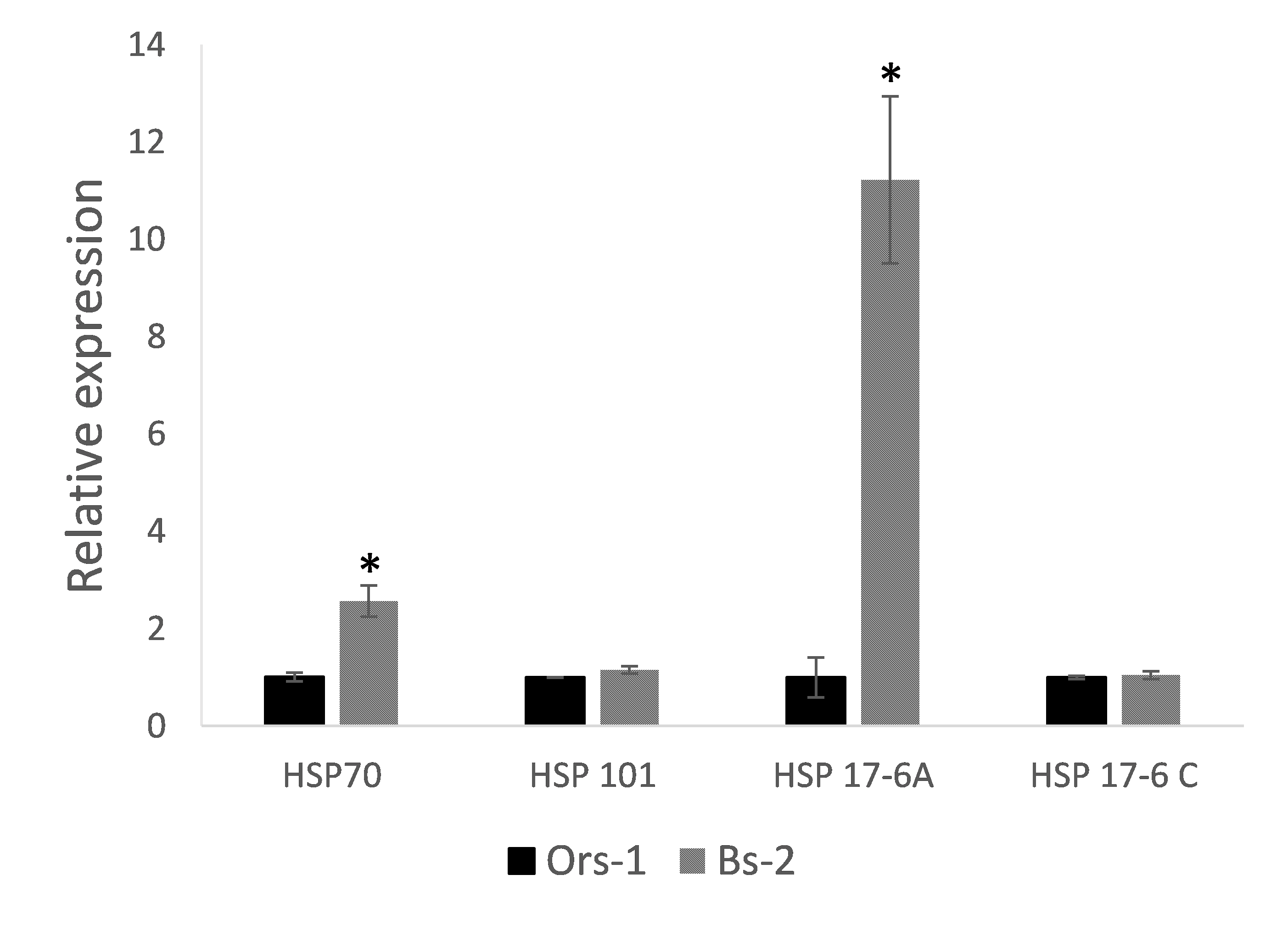

### Supplemental Figure 4

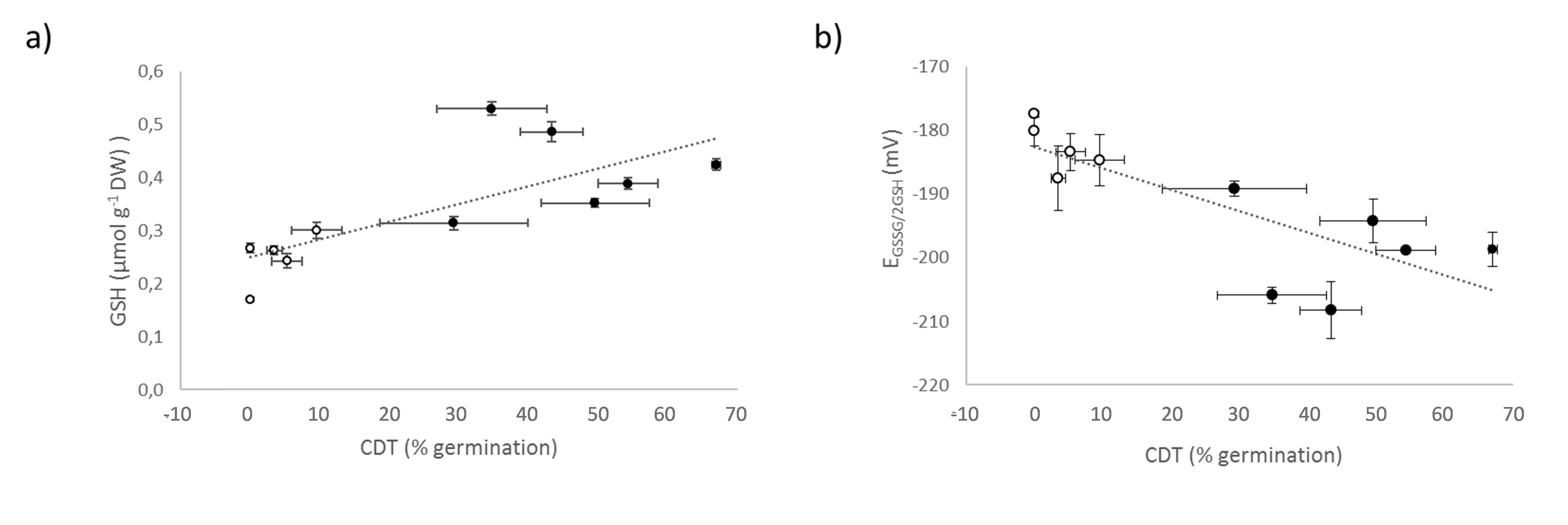
